# Shifting Beyond Classical Drug Synergy in Combinatorial Therapy through Solubility Alterations

**DOI:** 10.1101/2024.11.08.618644

**Authors:** Elham Gholizadeh, Ehsan Zangene, Uladzislau Vadadokhau, Danilo Ritz, Juho J. Miettinen, Rabah Soliymani, Marc Baumann, Mathias Wilhelm, Esko Kankuri, Paul A. Haynes, Caroline A. Heckman, Amir A. Saei, Mohieddin Jafari

**Affiliations:** Department of Biochemistry and Developmental Biology, Faculty of Medicine, University of Helsinki, Helsinki, Finland; Biozentrum, University of Basel, Basel, Switzerland; Institute for Molecular Medicine Finland (FIMM), Helsinki Institute of Life Science, iCAN Digital Cancer Medicine Flagship, University of Helsinki, Helsinki, Finland; Technical University of Munich, Munich, Germany; Department of Pharmacology, Faculty of Medicine, University of Helsinki, Helsinki, Finland; School of Natural Sciences, Macquarie University, North Ryde, New South Wales 2109, Australia; Department of Microbiology, Tumor and Cell Biology, Karolinska Institutet, Stockholm, Sweden

## Abstract

Acute myeloid leukemia (AML) remains a formidable clinical challenge due to genetic heterogeneity, high relapse rates, and toxicities associated with conventional chemotherapies. Rationally designed drug combinations offer improved efficacy, yet their selection is often empirical and lacks molecular mechanistic understanding. Here, we present CoPISA workflow (Proteome Integral Solubility/Stability Alteration Analysis for Combinations), a high-throughput proteomics workflow that captures protein solubility/stability alterations unique to combinatorial drug treatments, revealing mechanisms unattainable through single-drug analyses. Applying CoPISA to two rationally designed AML drug pairs, LY3009120-sapanisertib (LS) and ruxolitinib-ulixertinib (RU), we mapped primary (lysate) and secondary (living cell) protein target landscapes. Notably, our analysis uncovered an emergent mechanistic principle, “conjunctional targeting” (i.e., conjunctional inhibition), wherein cooperative drug actions induce treatment-specific targets not achievable individually, analogous to an AND-gate logic model. LS-specific AND-gate proteins converged on SUMOylation, chromatin condensation, and VEGF-linked adhesion, while RU-specific targets disrupted DNA-damage checkpoints, mitochondrial bioenergetics, and RNA-splicing machinery, collectively implicating synthetic-lethal vulnerabilities. Additionally, the post-translational modifications (PTMs) profiling of differential soluble proteins confirms several combination-induced modifications (e.g., acetylation, dimethylation, phosphorylation) on key AML proteins, such as *NPM1*. Network interrogation of AML-associated proteins showed that a high percentage of targeted proteins are unique to the combinations, including frequently mutated drivers *DNMT3A, NPM1,* and *TP53*. CoPISA exposes how drug pairs enact multi-axis pressure on AML cells through conjunctional targeting, a mechanistic layer beyond classical synergy. By pinpointing combination-exclusive protein targets and signaling pathways, CoPISA provides a blueprint for precision-guided regimen design in AML and other heterogeneous cancers. Data are available via ProteomeXchange with identifier PXD066812.

## Introduction

Acute myeloid leukemia (AML) is an aggressive hematologic malignancy characterized by the clonal expansion of immature myeloid cells in the bone marrow and peripheral blood, resulting in disrupted hematopoiesis ^1^. Due to the complexity of AML and the limited efficacy of monotherapies, combination therapy has long been central to treatment strategies ^2,3^. This approach often demonstrates superior efficacy compared to monotherapy, leveraging the synergistic or additive effects of different agents to combat complex diseases ^4^. However, challenges such as resistance to these combinatorial therapy regimes emphasize the need for further research to optimize treatment strategies and improve patient outcomes. On the other hand, most combinations to date have been derived empirically, with limited mechanistic understanding of how multi-agent treatments interact within the cellular proteome. This limitation is particularly problematic in AML, where genetic and epigenetic heterogeneity fosters divergent responses to therapy and promotes the rapid emergence of resistant clones ^5^. Moreover, high toxicity associated with multi-drug regimens restricts their broader clinical application and often necessitates dose reductions that compromise efficacy ^6,7^.

Proteomics-based approaches have recently emerged as promising tools for elucidating drug mechanisms of action at the systems level. One such approach is the study of drug-induced shifts in protein conformational solubility/stability, which can provide insights into the potential protein targets and then mechanisms of action of therapeutics ^8,9^. The thermal shift assays leverage the principle that drug binding affects the solubility or thermodynamic stability of a protein, allowing researchers to identify drug targets across the whole cellular proteome^10^. In this line, the PISA assay is a high-throughput technique to monitor drug target engagement and has been employed to decipher the targets of hundreds of drugs recently ^10,11,12^. However, these methods have been primarily applied to single-agent treatments and fail to capture emergent molecular effects unique to drug combinations. To overcome this gap, we developed CoPISA (Proteome Integral Solubility/Stability Alteration Analysis for Combinations), a high-throughput proteomics workflow designed to interrogate combination-specific solubility alterations in proteins.

The CoPISA method captures drug interactions by detecting changes in protein solubility/stability that occur specifically when a combination of drugs is used, as compared to single-drug and control samples. We applied this approach to two recently identified drug combinations with demonstrated high efficacy and low toxicity in AML patient samples ^13^. This technique uses advanced proteomic tools to explore both primary (using lysate sample) and secondary (using living sample) target effects, providing a comprehensive understanding of how combinatorial therapies can modulate multiple signaling pathways to overcome treatment resistance ^10,14^. This research extends beyond traditional drug target identification by examining how these interactions impact cellular behavior at a systems level, thereby addressing current limitations in understanding the mechanisms behind successful drug combinations. Moreover, we introduce a molecular mechanism termed ‘conjunctional targeting,’ where the combined action of drugs produces biological responses that are unattainable by individual compounds (more definition in the Result section). This perspective transforms our understanding of drug interactions and paves the way for more effective therapeutic strategies in AML and other complex diseases.

## Results

### CoPISA Assay principal and *Ab Initio* Simulation

To investigate how combinatorial drug treatments affect protein solubility, we used the CoPISA workflow, an extension of the PISA assay tailored for combinatorial therapies. In the CoPISA assay, we quantify protein solubility/stability by measuring the total amount of each protein, represented as the area under the melting curve of each protein (*S*_m_), enabling precise solubility/stability measurements without the need for curve fitting. The CoPISA workflow involves treating cells with each drug individually (e.g., A and B), as well as in combination (AB), alongside untreated controls. Lysates or living cells are then subjected to thermal treatment across a temperature gradient to assess shifts in the amount of soluble/stable proteins (*S*_m_) under each condition (**Fig. 1**). Relative changes in *S*_m_ compared to the control (Δ*S*_m_) provided quantitative insight into drug-induced effects on protein solubility/stability. These shifts indicate either stabilization (increased *S*_m_) or destabilization (decreased *S*_m_) of proteins. Note that in the following, we used “solubility” shift instead of “solubility/stability” to emphasize the change in the amount of soluble protein after altering stability with increasing temperature in the presence and absence of drugs.

**Figure 1:**
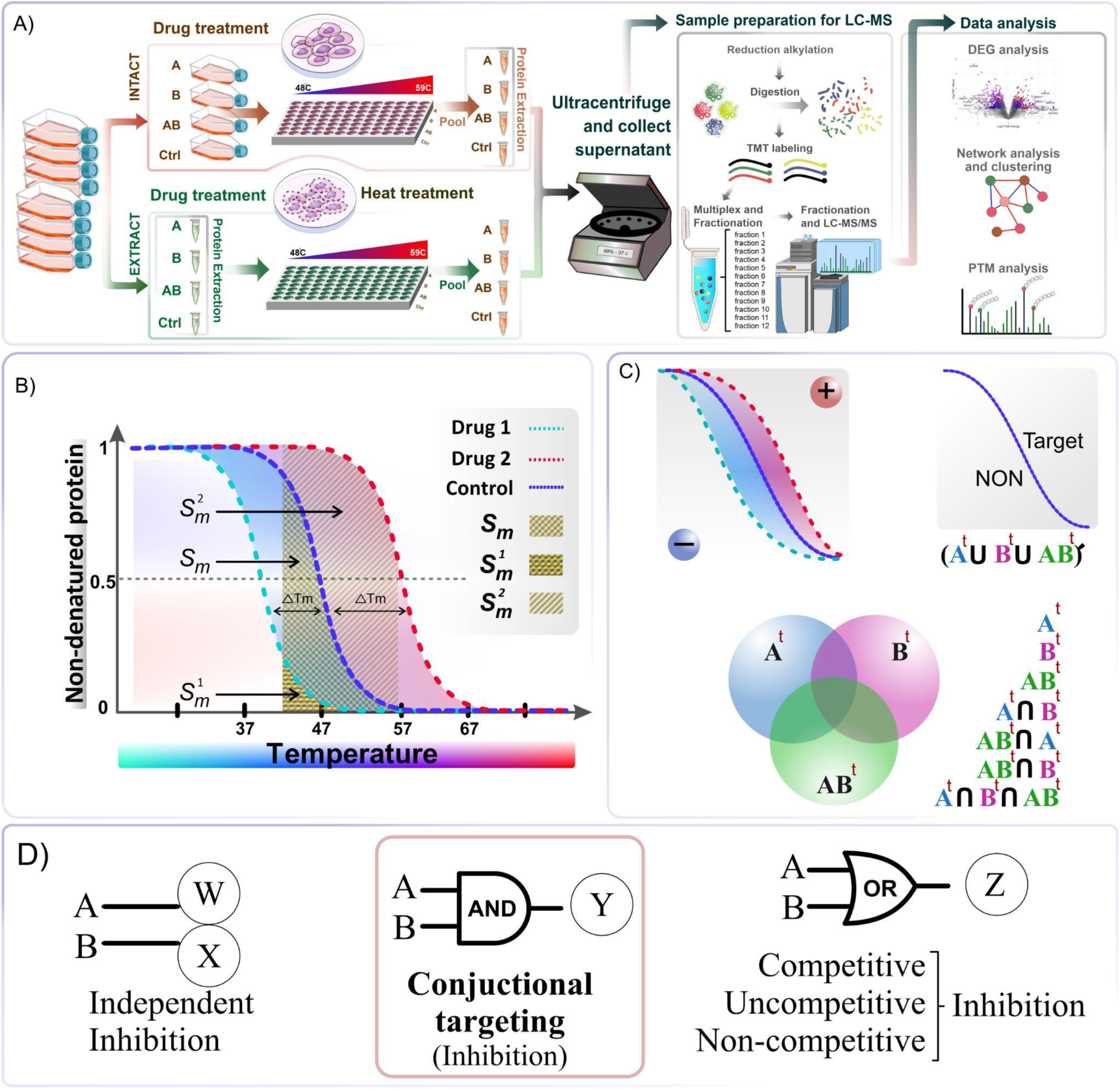
CoPISA workflow and possible states for target discovery in combinatorial therapy. (A) Schematic overview of the CoPISA method applied to both intact cells and lysates. Cells are treated with Drug A, Drug B, or their combination (AB) and control, followed by heat treatment across a temperature gradient (48–59°C). Soluble proteins are isolated post-ultracentrifugation and analyzed by LC-MS, with subsequent downstream analyses. (B) Comparison of the PISA concept of integral analysis with the concept of sigmoidal curve fitting when using two drugs. In the PISA workflow, protein solubility shifts are measured based on the area under the curve (Δ*S_m_*) rather than using the median of the fitted sigmoidal curve of denatured protein (Δ*T_m_*) at several temperature points. The area under the curve for the first drug (light blue curve, *S_m_^1^*) and the second drug (red curve, *S^2^_m_*) is used to calculate Δ*S* within the temperature range of 48–59°C. The same way is used to measure the area under the curve for the combination treatment. (C) Theoretical illustration of the CoPISA concept. This panel illustrates comparisons across multiple single treatments, including drug A, drug B, and combination AB, and various intersections of identified potential targets (i.e., *A^t^*, *B^t^*, *AB^t^*) of these three treatments compared to control (e.g., *AB^t^*∩*A^t^*, *A^t^*∩*B^t^*, *A^t^*∩*B^t^*∩*AB^t^*, *AB^t^*∩*B^t^*). It also highlights cases where no target exists for the union of different treatments (*A^t^* ∪ *B^t^* ∪ *AB^t^*). (D) Mapping of drug inhibition models to logical gate analogies: Independent inhibition, where drugs A and B act separately on targets W and X, respectively, corresponds to an independent action; conjunctional targeting, where drugs A and B jointly affect a single target Y, aligns with the *AND* logical gate; and *competitive*, *uncompetitive*, and *non-competitive inhibition*, representing mechanisms that are either mutually exclusive or involve overlapping inhibition effects, correspond to the *OR* logical gate.

The CoPISA workflow enables a detailed analysis of drug-target interactions by classifying proteins into eight classes, revealing distinct impacts on protein solubility. The four trivial classes of proteins according to drug-target interactions can be hypothesized as follows: A (protein targeted by drug A), B (protein targeted by drug B), A∩B (protein targeted by both drug A and drug B), A⋃B (protein not targeted by any drug) when using two drugs or compounds for treatment. In one class, some proteins are unaffected by all treatments, showing no changes in solubility compared to the control. Two classes involve *independent inhibition* (A or B), where treatment with either drug A or drug B alone leads to specific alterations in solubility profiles, indicating individual effects on protein solubility. These proteins could suggest a classical synergy mechanism in combinations by targeting two proteins from different or identical biochemical pathways. A∩B represents proteins targeted by both drug A and drug B, where combined treatment results in a distinct solubility shift, potentially indicating cooperative or enhanced binding effects. These proteins could highlight synergistic interactions where the combined action of both drugs leads to a stronger impact on solubility than either drug alone.

As a theoretical proof of principle, we conducted a simulation of melting sigmoidal curves for 10,000 hypothetical proteins to explore the possibility of all four classes. This preliminary computational analysis involved randomly selecting melting temperatures *T*_m_ in the range from 48 to 56°C using error (erf) and square root (sqrt) functions. Examples of thus-simulated melting curves are shown in **Fig. 2**, illustrating the variation in thermal stability across the different protein classes. Our analysis suggests that all four expected classes of targeting proteins for this two-drug combination are theoretically possible, based on a normal distribution and hypothetical values for the mean and variance (**Fig. 2A**). This finding supports the idea that different classes of proteins could be differentially affected by the drug combination. In addition, we observed a similar pattern of these four classes with slightly distinct distribution based on the mean and variance values extracted from a proof-of-principle TPP published article ^15^ (**Fig. 2B**), further validating the robustness of our theoretical model and aligning with previously observed experimental data.

**Figure 2:**
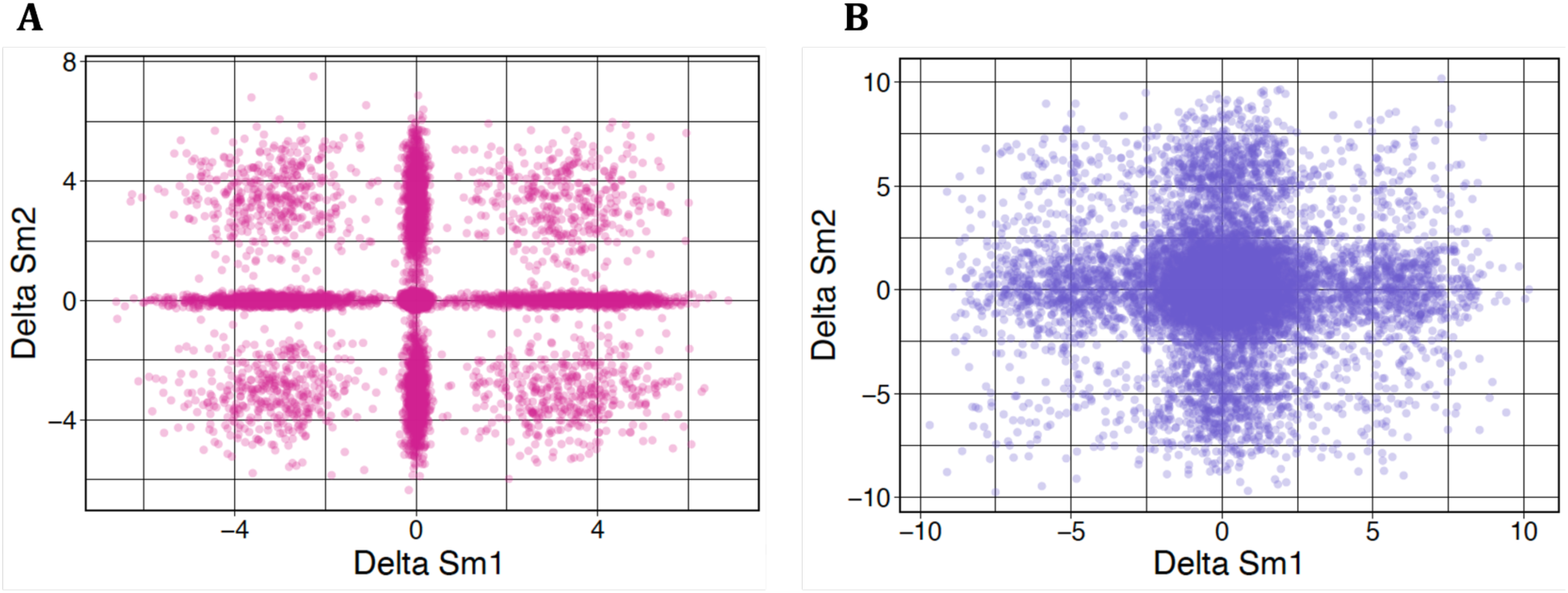
The simulated relationship of protein thermal shift of two hypothetical drugs. **(A)** Δ*S^1^* and Δ*S^2^* in 10,000 proteins were simulated and depicted using a normal distribution with hypothetical mean and variance, and (B) with mean and variance extracted from the proof-of-principle TPP published article^15^.

However, in the CoPISA workflow, we also consider the treatment of the combination (AB) in addition to the individual drugs. This introduces a unique condition where certain proteins are exclusively targeted by the combination treatment (AB), requiring the simultaneous presence of both drugs to induce significant shifts in solubility. We refer to this as *conjunctional targeting (i.e., conjunctional inhibition)*. This cooperative effect can be framed within logical gate theory^16^, suggesting an *AND* gate mechanism; both drugs must be present to induce inhibition sufficiently. This pattern indicates that these proteins may possess multiple binding sites or complex allosteric regulation modulated only by both drugs, providing insights into the underlying mechanisms of this drug combination. Conversely, if a protein is targeted in the presence of A, B, or A∩B, it aligns with an *OR* gate model, representing an overlap of effects across treatments. This mechanism offers an additional explanation for synergy in drug combinations, where the effects are enhanced when used in tandem compared to monotherapy.

Ultimately, we identified eight classes of proteins. Four of these classes demonstrate shared protein targets across all drug treatments (*AB^t^***∩***A^t^*, *A^t^***∩***B^t^*, *A^t^***∩***B^t^***∩***AB^t^*, *AB^t^***∩***B^t^***), w**hile two classes show individual protein targets specific to drug A or B treatments. One class, termed *conjunctional targeting*, is uniquely targeted by the combination treatment (AB), and the remaining class consists of proteins that are not targeted by any treatment.

### Global CoPISA Profiling and Intersectional Target Analysis in AML Cell Lines

We applied the CoPISA workflow to four AML cell lines, MOLM-13, MOLM-16, SKM-1, and NOMO-1, to examine how each single agent (LY3009120, sapanisertib, ruxolitinib, and ulixertinib) and its corresponding combination (LS and RU) reshape the soluble proteome. Both lysates (capturing direct, primary binding events) and intact cells (capturing the net effect of primary and secondary interactions) were analyzed **(Supplementary File S1)**.

Volcano plots (Fig. 3A**–D**) illustrate a striking yet therapy-specific redistribution of protein solubility. In the living cell condition (Fig. 3A), treatment with ruxolitinib, ulixertinib, and the RU combination resulted in 334, 393, and 553 significantly altered proteins, respectively. In the lysate (Fig. 3B), treatment with ruxolitinib, ulixertinib, and the RU combination yielded 350, 479, and 146 significantly altered proteins, respectively. Corresponding to intact or living cells (Fig. 3C), treatments with LY3009120, sapanisertib, and the LS combination yielded 349, 293, and 308 significantly altered proteins, respectively. The cell lysates treated with LY3009120, sapanisertib, and the LS combination resulted in 270, 141, and 215 significantly altered proteins, respectively (Fig. 3D). These findings highlight that, while individual treatments tend to produce extensive alterations in protein solubility, the combination treatment exhibits a more selective pattern of solubility changes, particularly under the lysate condition. Together, these observations underscore the differential impact of single and combined treatments on protein solubility, with notable differences between lysate and living cell conditions that may inform the distinct mechanistic effects of combinatorial therapies on protein dynamics.

**Figure 3.**
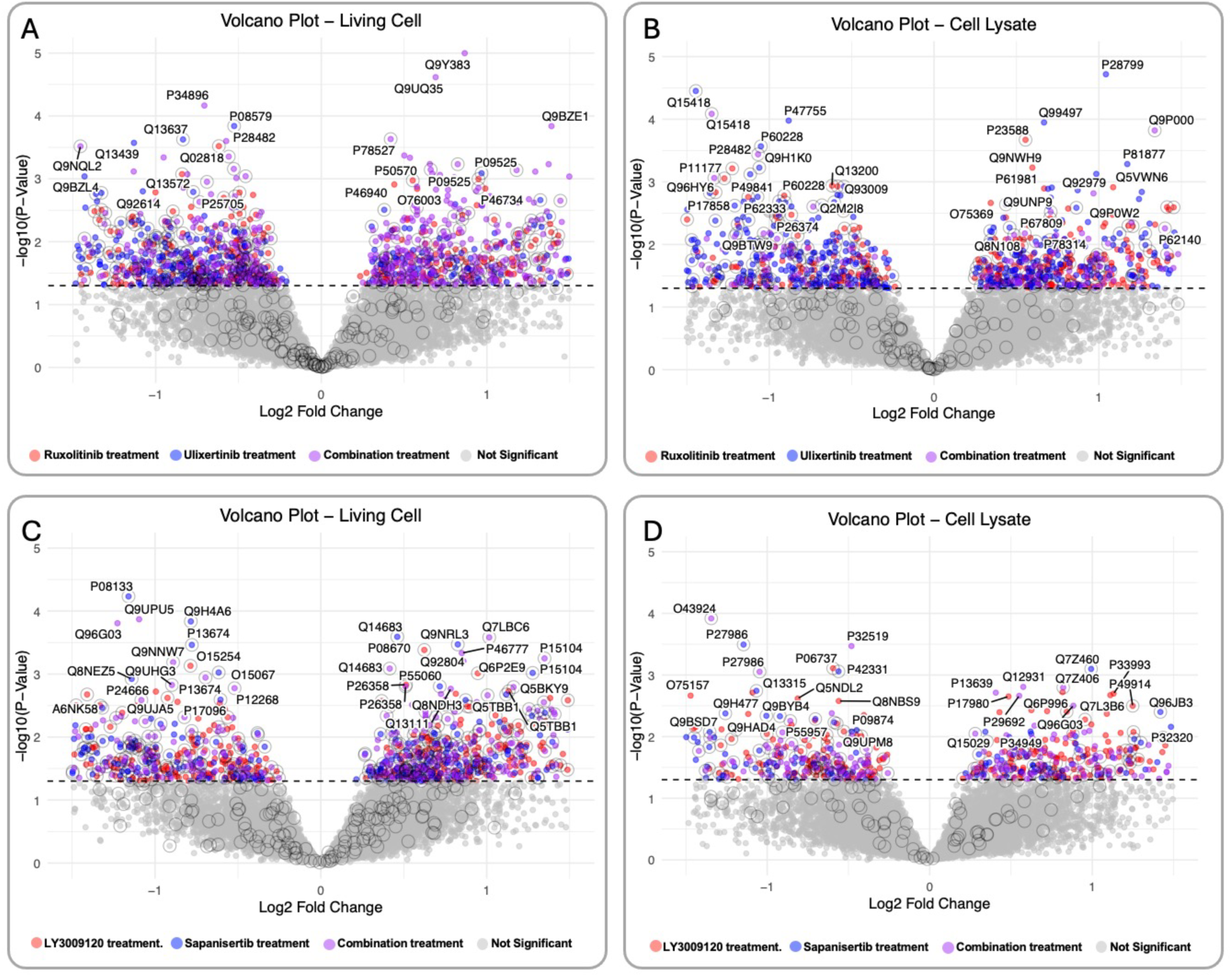
Volcano plots reveal protein targets identified by the CoPISA method in lysate and living cell treatments with drug combinations. Each panel is a superimposition of three comparisons between control and treatment groups, with solubility shifts indicated by different colors: living cells (A) and lysate cell samples (B) treated with ruxolitinib (3 uM) (red), ulixertinib (3 uM) (blue), and their combination, RU (purple); and living cells (C) and lysate cell samples (D) treated with LY3009120 (500 nM) (red), sapanisertib (500 nM) (blue), and their combination, LS (purple). Dots with outlines represent proteins uniquely identified in the corresponding treatment.

To ensure the robustness of the significance cutoff and to effectively control the false discovery rate (FDR) for solubility shifts, a comprehensive permutation analysis of protein S_m_ values was conducted. The analysis demonstrated that, on average, the FDR across all twelve comparison analyses was maintained at 5.25% for 12000 permutations, ensuring the reliability of the identified significant solubility shifts. Taken together, these data show that each regimen reshapes the AML proteome in a unique and context-dependent manner.

To determine how consistently the two drug pairs engage their intracellular targets, we compared proteins whose solubility was significantly altered in lysate-based (primary targets) and intact-cell (primary + secondary targets) CoPISA experiments (Fig. 4). Across all treatments, the living-cell fractions yielded more hits than lysate samples, as expected from the metabolic activity of intact cells and the capture of both primary and secondary protein targets. Notably, the LY3009120– sapanisertib (LS) combination shifted the solubility of 177 proteins in living cells but only 147 in lysate; ruxolitinib–ulixertinib (RU) affected 414 and 89 proteins, respectively. These partially overlapping target sets point to distinct intracellular consequences of the two regimens.

**Figure 4:**
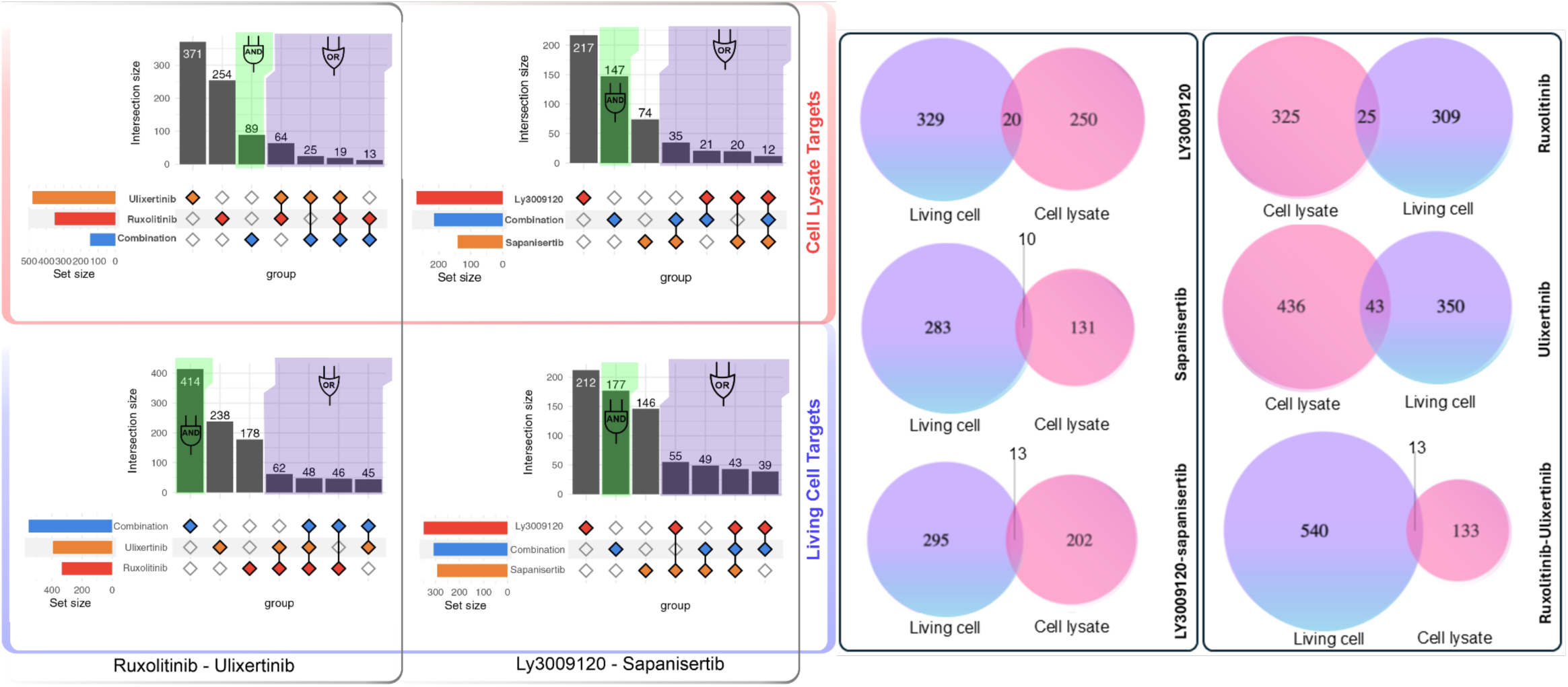
Comparison of protein targets identified in lysates and living cells across drug treatments. UpSet plots show the number of protein targets identified in lysate and living cell samples following treatment with LY3009120–sapanisertib and ruxolitinib–ulixertinib. Venn diagrams further illustrate the distribution of targets detected using the CoPISA method across both sample types—living cells and lysates—after treatment with individual drugs and their combinations. In each diagram, targets unique to living cells are shown in blue, those unique to lysates in pink, and shared targets in the purple overlap region. These visualizations provide insights into drug-dependent protein interactions under different cellular conditions.

In Fig. 4, the CoPISA approach was used to compare drug-dependent target engagement between living cells and cell lysates. Each Venn diagram shows the distribution of targets unique to living cells, unique to lysates, and shared between both. While most targets were condition-specific, a notable number of overlapping proteins, particularly in the Ruxolitinib (25), Ulixertinib (43), and combination treatments (13 each), serve as a strong validation of the CoPISA method across these distinct experimental conditions. These shared targets reflect consistent drug-protein interactions detectable in both intact and lysed cellular environments, supporting the robustness of CoPISA in mapping solubility shifts associated with drug binding.

In samples treated with LY3009120, 20 proteins with significant solubility shifts are consistently detected in both living cells and cell lysate samples (Fig. 4). Among these, PTPN11, encoding the SHP2 phosphatase frequently mutated in myeloid malignancies, suggests inhibition of RAS/MAPK pathway signaling critical for leukemic cell maintenance ^17^. In sapanisertib-treated samples, 10 proteins are shared between lysate and intact cell conditions. Notably, S100A8, a calcium-binding protein implicated in inflammatory signaling and leukemogenesis suggests a role for sapanisertib in modulating immune-related pathways ^18^. Other intersected proteins, such as UCHL1 and TPM1, suggest modulation of protein turnover and cytoskeletal dynamics, respectively, which may influence AML cell viability ^19,20^. Under the combination-treated samples, 13 proteins are shared in lysed and intact samples, while under the AND gate, seven proteins are targeted in samples treated with the combination of LS in both living cell and lysate samples. Among these, ICAM3, an adhesion molecule, suggests effects on leukemic cell adhesion and trafficking within the bone marrow niche ^21^. Collectively, these findings support the hypothesis that the LS combination elicits conjunctional targeting through cooperative targeting of signaling, adhesion, and trafficking mechanisms.

In the ruxolitinib-treated samples, 25 proteins are targeted in both living cell and lysate samples (Fig. 4). Of particular note, ABL1, a non-receptor tyrosine kinase, plays critical roles in leukemogenesis and resistance to therapy, suggesting that ruxolitinib may suppress oncogenic kinase signaling in AML ^22^. Similarly, in ulixertinib-treated samples, 45 proteins were shared between lysate and living cell conditions. The detection of MAPK3 (ERK1), a central node in the MAPK pathway, underscores inhibition of proliferative signaling cascades by ulixertinib ^23^. Under the combination treatment, 13 proteins are shared in lysate and intact samples, and four proteins are specific to the combination and are not targeted in single treatment. TTC19, a mitochondrial protein involved in respiratory chain complex III assembly, suggests that RU combinatorial therapy may compromise mitochondrial bioenergetics, leading to enhanced apoptotic priming of AML cells ^24^. These conjunctional targets reinforce the concept that RU combinatorial therapy reprograms key cellular processes to induce synthetic lethality in leukemic cells.

Taken together, these updated intersectional analyses provide deeper insight into how single and combination treatments differentially engage cellular targets in AML, with the AND-gate proteins emerging as critical effectors of combinatorial synergy. These findings highlight therapeutic vulnerabilities that could inform the rational design of precision-guided combinatorial regimens.

To further investigate the association patterns in protein solubility alterations induced by combination treatments, we generated fold change-fold change (FCFC) plots (Fig. 5). These plots compare the solubility alterations in proteins for each combination treatment relative to the effects observed with the individual drugs alone. Specifically, the FC-FC analysis examines (1) combination vs. drug 1 (e.g., LS combination vs. LY3009120 alone) and (2) combination vs. drug 2 (e.g., LS combination vs. sapanisertib alone). Interestingly, only 0.84% of protein alterations exhibit inconsistent directional changes, where stabilization or destabilization patterns differ between combination and single-drug treatments. This finding highlights a strong alignment between the solubility alteration patterns induced by the combination treatments and those observed for individual drugs, particularly from a stabilization perspective. These results reinforce the idea that combination treatments largely maintain the stabilization directionality of the single drugs, providing further evidence of their targeted and coordinated action on protein solubility.

**Figure 5:**
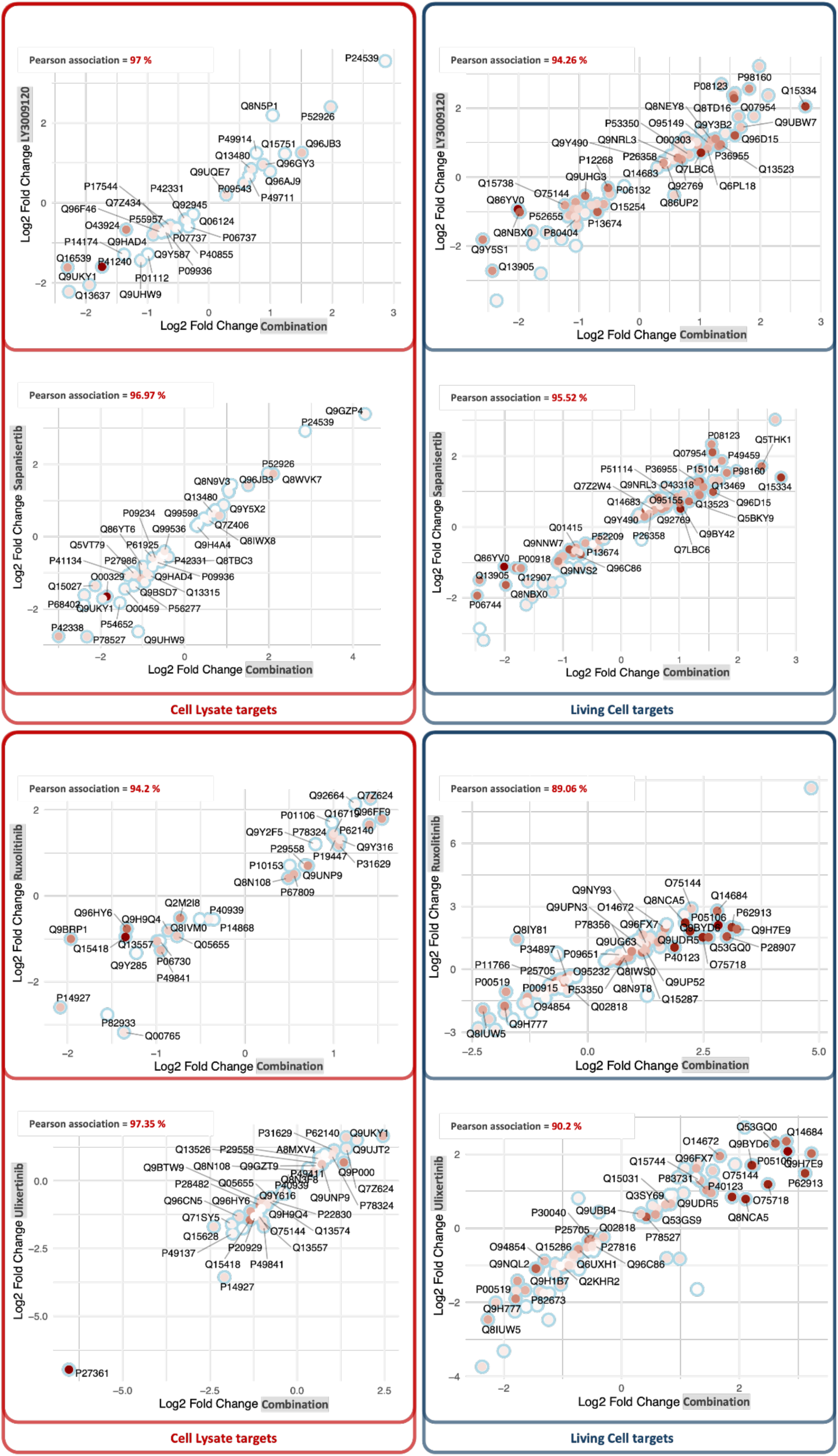
Pairwise fold change comparisons of treatment conditions based on significant protein targets. The FC-FC plots display common protein target IDs with fold change values for each treatment relative to control samples. These plots are presented separately for cell lysate (red frame left panels) and living cells (dark blue frame right panels), with distinct panels for each combination.

### Exploring Drug-Drug Interactions Through Mass Spectrometry Analysis

In the next phase, potential noncovalent interactions or unforeseen reactions within the selected drug combinations were examined to ensure that the observed effects on protein solubility were independent and specific to each compound. Using LC-MS/MS analysis^25^, we compared the spectral profiles of each drug alone with those of the combined samples (Fig. 6). By analyzing the MS1 spectra and extracted ion chromatograms (XICs) of each compound separately and in combination, we assessed whether a new profile emerges from reacting or interacting drug molecules.

**Figure 6:**
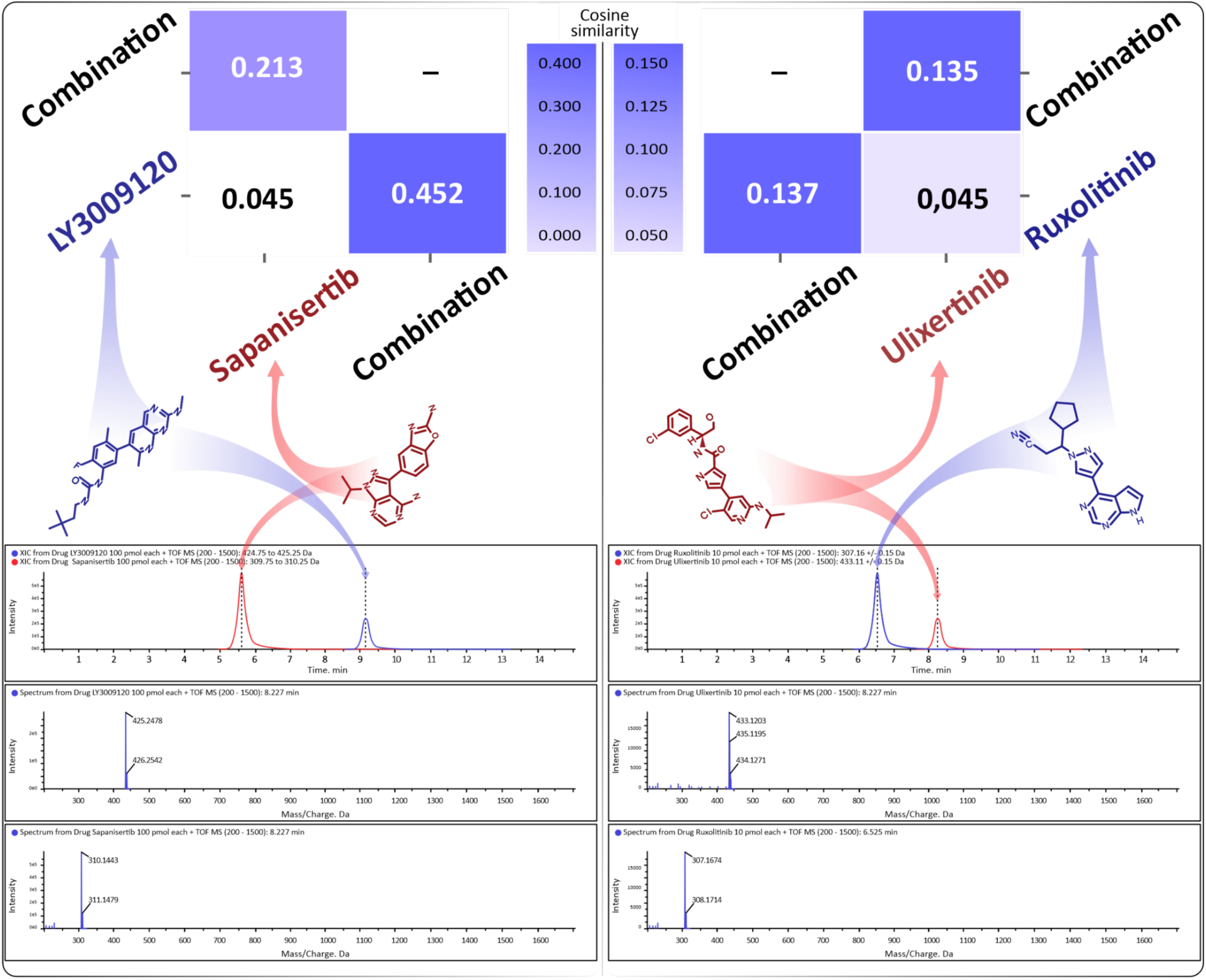
Mass spectrometry analysis of four individual compounds and their combinations. Molecular structures of each compound are shown alongside their MS1 spectra and total ion currents (TICs) for the combined samples. For each drug combination, TIC plots are provided to display the corresponding chromatographic profiles and detected mass-to-charge (m/z) values for each compound. Pairwise cosine similarity of TICs between individual compounds and their combinations is presented based on quantified MS peak intensities.

The XICs and MS1 spectra of the combined drugs showed no other peaks or mass shifts other than initial single-drug ion peaks, indicating an absence of detectable interactions or reactions between the compounds or their modification. Additionally, cosine similarity analysis of the whole MS1 spectra provided a comparative view of each drug combination. This analysis confirmed that the effects observed on protein targets were attributable to specific mechanisms of each drug, rather than nonspecific interactions within the combinations. Each compound displayed stable peak intensities and retention times across conditions.

These findings support the hypothesis that the protein targets identified for the LS and RU combinations arise from the concurrent, yet independent, actions of each drug, rather than from non-specific interactions between them. This reinforces the proposed “AND gate” inhibition mechanism in combinatorial therapy, suggesting these combinations achieve synergistic effects through coordinated drug action and potential engagement of new signaling pathways.

### Mechanistic Insights into “AND” and “OR” Gate Targeting Patterns

To gain deeper insight into the mechanisms underlying the drug combinations, we performed comparative pathway enrichment analysis of proteins exhibiting significant solubility shifts in response to LS and RU treatments. These analyses leveraged both lysate and intact cell fractions to distinguish context-specific biological processes perturbed under combinatorial treatment. The framework of “AND” and “OR” logic gates allowed us to differentiate between synergistic effects of combinatorial therapies, i.e., independent or conjunctional targeting (Fig. 7**, Supplementary File S2)**. This systems-level approach uncovered molecular signatures that converge on key vulnerabilities in AML biology.

**Figure 7.**
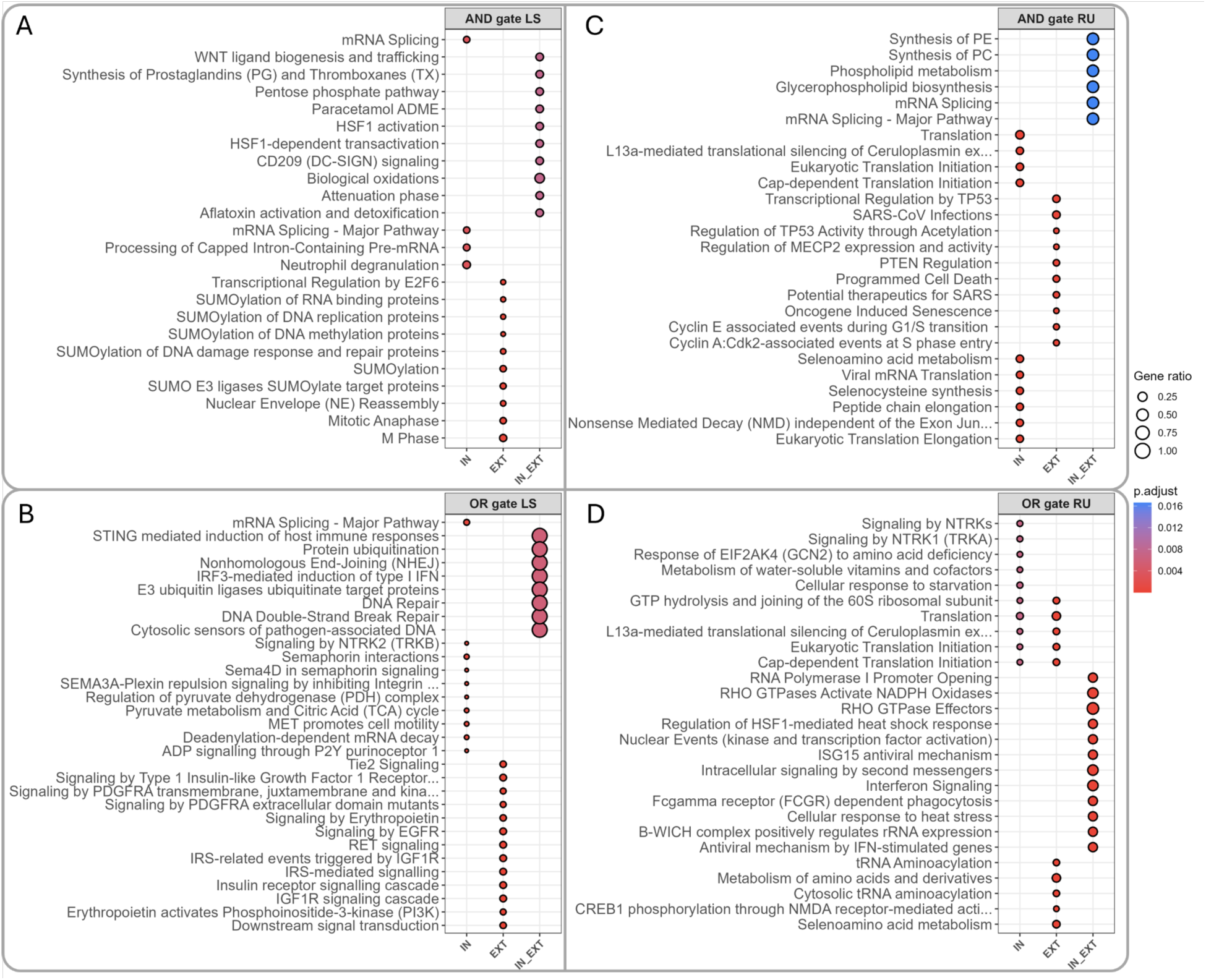
Bubble plots illustrating the top 15 enriched Reactome pathways (p.adj < 0.05) for each combination and gating logic. If no pathways meet the p.adj < 0.05 threshold, the top 10 pathways ranked by adjusted p.adj are shown. Facets display AND-gate (conjunction) and OR-gate (disjunction) comparisons for LY3009120 & sapanisertib and ruxolitinib & ulixertinib. Within each panel, bubbles at “IN” represent pathways unique to intact (live) samples, “EXT” indicates pathways unique to extract (lysate) samples, and “IN_EXT” identifies pathways common to both. Bubble size corresponds to gene ratio (pathway genes / total tested), and color intensity reflects raw p-values (red = lower, blue = higher). Pathway names are truncated for clarity.

Within the “AND” gate, representing proteins selectively responsive to the LS combination but not either single agent, significant enrichment is observed in SUMOylation processes as primary targets (Fig. 7A). This points to disruption of genome stability and transcriptional control pathways, which are critical for leukemic maintenance and progression ^26^. Enrichment of mitotic phase processes, including chromatin condensation and nuclear envelope dynamics, further indicates interference with cell division fidelity, a hallmark vulnerability in rapidly proliferating AML cells ^27^. Conversely, enrichment analysis of proteins from living cell samples, which represents downstream pathways beside direct targets, highlights RNA processing and immune function. These findings suggest that the combination treatment predominantly interferes with transcriptional output and modifies immune interactions within the AML microenvironment ^28^. Proteins shared across lysate and living cell fractions under the “AND” gate configuration reveal additional points of vulnerability. These include activation of stress-responsive transcription, redox homeostasis, metabolic rewiring, lipid signaling, and immune remodeling. Disruption of WNT signaling and CD209-mediated pathways further implicates LS in undermining both developmental and innate immune axes, processes increasingly linked to AML cell persistence and therapy resistance ^29,30^. These converging effects underscore a multi-axis destabilization of AML cell resilience.

Further analysis of the “OR gate” under LS treatment reveals distinct yet interconnected patterns between lysate and living cells (Fig. 7B). Lysate-specific proteins prominently involve mitogenic and growth factor signaling cascades, particularly receptor tyrosine kinase-driven (RTKs) pathways, emphasizing interference with key proliferative and metabolic regulatory circuits. Targeting these pathways is crucial, as RTK-driven signaling often sustains leukemic growth and survival, making their inhibition a promising therapeutic approach in AML ^31^. In contrast, living cell-specific pathways are heavily enriched in translation and protein homeostasis mechanisms, implicating significant disruptions to ribosome assembly, protein synthesis, and mRNA surveillance. This reflects suppression of biosynthetic output and stress-induced translational reprogramming, which AML cells rely on to sustain high proliferative demands and evade treatment stress ^32^. Importantly, shared targets between both fractions, although fewer in number, converge on critical biosynthetic processes, including phospholipid metabolism and RNA splicing. These pathways govern membrane integrity and transcriptome regulation, and their coordinated disruption may destabilize intracellular architecture and gene expression fidelity. Altogether, the OR-gate analysis reveals how LS imposes widespread molecular pressure by targeting essential survival signaling, metabolic infrastructure, and RNA regulatory mechanisms, driving cumulative cellular stress that likely exceeds the adaptive capacity of AML cells.

In parallel, the RU combination demonstrates complementary mechanistic patterns (Fig. 7C). The “AND” gate analysis for RU identifies lysate-specific enrichment in transcriptional regulation, cell cycle control, apoptosis, and epigenetic modulation, which reflect primary disruption of proliferative signaling and tumor suppressor networks frequently dysregulated in AML ^33^. Living cell-specific pathways emphasize translation control and protein homeostasis, indicating significant disruptions to ribosome assembly, protein synthesis, and mRNA surveillance, pathways crucial for maintaining the high biosynthetic and metabolic demands of AML cells ^34^. Shared targets in both fractions consistently impact phospholipid metabolism and RNA splicing, underscoring a unified effect on fundamental biosynthetic processes fundamental for AML cell integrity and proliferation ^35^. These findings highlight the ability of RU to inhibit biosynthetic machinery critical for AML cell survival and proliferation.

Similarly, OR-gate analysis of RU drug combination treatment across lysate and intact cell fractions identifies extensive disruptions in translational control and stress response pathways that are critically relevant to AML pathobiology (Fig. 7D). Lysate samples emphasize modulation of protein biosynthesis, amino acid metabolism, and transcriptional regulation, core processes that are often upregulated in AML to support unchecked proliferation and metabolic adaptation ^30^. In the intact cells, RU highlights interference with translation initiation and metabolic stress sensing, which are essential for AML cells to withstand nutrient limitation and therapeutic pressure ^36^. Shared proteins underscore activation of innate immunity and redox regulation, processes that are increasingly recognized for their role in AML immune evasion and oxidative stress tolerance ^29^. These convergent mechanisms suggest that RU therapy disrupts not only biosynthetic and metabolic robustness but also immunomodulatory axes, delivering a comprehensive blockade of AML cellular resilience and survival.

In conclusion, comparative “AND” and “OR” gate analyses highlight how LS and RU combinations distinctly yet convergently target AML vulnerabilities. LS primarily disrupts proliferative signaling and metabolic regulation, with secondary effects on RNA processing and immune modulation. Shared vulnerabilities under LS include phospholipid metabolism, redox balance, and RNA splicing, collectively destabilizing AML cell survival networks. Similarly, RU primarily impairs translational control and stress response pathways, with secondary disruption of immune regulation and biosynthetic processes. Shared targets for RU converge on phospholipid metabolism and RNA processing, undermining AML biosynthetic and adaptive capacity. These findings illuminate critical molecular nodes and provide a mechanistic rationale for designing combinatorial therapies to overcome resistance and improve AML outcomes.

### Roles of CoPISA-Identified Protein Targets in AML-Associated Signaling Networks

We performed a comprehensive functional interaction network analysis of AML-associated proteins to better understand the mechanistic implications of the combinatorial therapies LY3009120-sapanisertib (LS) and ruxolitinib-ulixertinib (RU). The AML-associated network is constructed by integrating Reactome signaling interaction data with high-confidence AML-associated genes derived from DepMap ^37^ and the mutation dataset of the FIMM-AML cohort study (p < 0.001, high-impact variants) ^38^. Directed shortest paths connecting AML-associated protein pairs result in a robust subnetwork containing interconnected components reflective of critical AML biology.

To distinguish treatment-specific perturbations within the AML network, we quantified the overlaps of AML subnetwork genes modulated by each combinatorial regimen **(Supplementary** Figure 1). This comparative analysis reveals that LS treatment modulates a total of 34 AML-associated genes, with 16 genes (47%) uniquely targeted by this combination and not by either single drug. This relatively high proportion of combination-specific targets underscores the synergistic potential of LS to engage molecular vulnerabilities that remain inaccessible to monotherapies. Similarly, RU treatment perturbs 37 AML-associated genes overall, of which 20 genes (54%) are distinct to RU, further highlighting the capacity of this combination to reprogram AML-specific networks in a manner that extends beyond additive effects of its individual components. These findings indicate that both LS and RU elicit a substantial degree of unique network rewiring, suggesting they engage complementary biological processes and may overcome redundant survival pathways exploited by leukemic cells under single-agent treatment.

Building on this annotated, treatment-layered AML subnetwork, we applied community detection, identifying 48 discrete clusters. Eleven clusters (i.e., C1, C2, C3, C4, C5, C6, C7, C8, C9, C10, and C16), prioritized based on a size threshold (≥10 genes) and overlap with CoPISA solubility shifts, were annotated using Reactome over-representation analysis (FDR ≤ 0.05). These clusters represent functional modules ranging from epigenetic regulation to immune signaling, offering a systems-level view of therapy-induced vulnerabilities (Fig. 8**, supplementary File S3)**.

**Figure 8.**
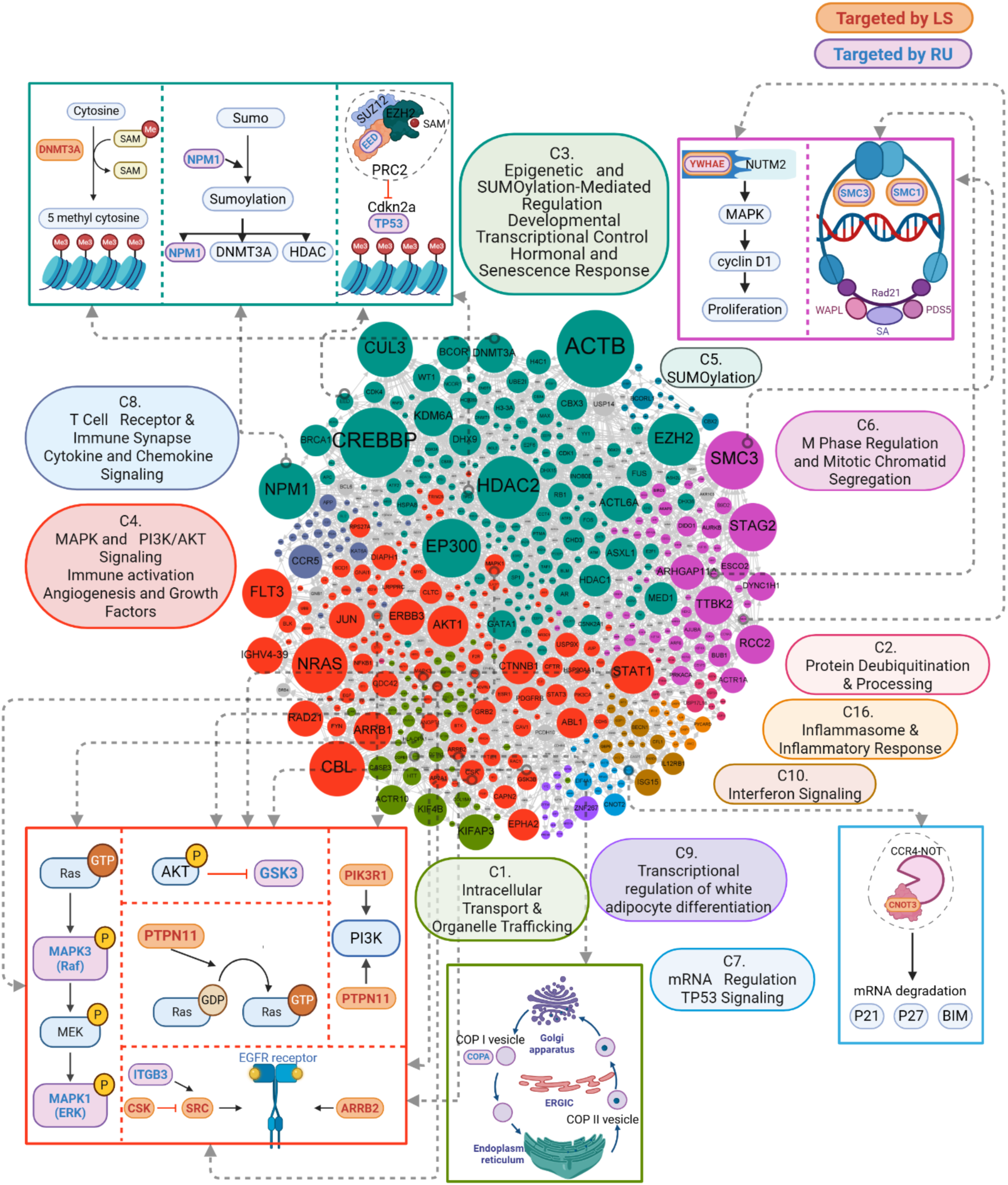
Functional interaction network of AML-associated genes revealing key proteins and pathways targeted by LS and RU combinatorial therapies. The network was constructed by integrating Reactome signaling interactions with high-confidence AML-associated genes derived from DepMap and the mutation data of the FIMM-AML cohort study (*p* < 0.001, high-impact variants). Directed shortest paths between all AML-associated protein pairs were used to define an induced subnetwork. Community detection using the Walktrap algorithm identified 48 discrete clusters, of which 11 are visualized in this plot. Each cluster is annotated via Reactome pathway enrichment (*adjusted p* < 0.05), highlighting distinct functional modules. Proteins significantly affected by combination treatments, LS (orange) and RU (purple), are annotated based on CoPISA solubility shifts, highlighting treatment-specific network perturbations. Peripheral schematics, framed using the same color codes as their corresponding clusters, highlight key pathways and molecular mechanisms enriched within major clusters, linking proteomic alterations to underlying biological processes.

The prioritized clusters highlight distinct processes significantly enriched within the AML network context, reflecting both shared and unique vulnerabilities to LS and RU treatment. Cluster 1 (C1), enriched for intracellular transport and organelle trafficking, shows significant disruption in COPI-mediated Golgi-ER transport pathways, critical processes for leukemic cell viability. These findings align with evidence linking lysosomal trafficking to AML survival, metabolism, and therapy response, highlighting the therapeutic potential of targeting cellular logistics ^23, 39^. Clusters 10, 16, and 8 are enriched in immune and inflammatory signaling, including interferon, inflammasome activation, and cytokine pathways, underscoring the central role of immune modulation in AML pathogenesis and treatment response.

Cluster 3 (C3) is strongly associated with epigenetic regulation, chromatin remodeling, SUMOylation, and PTEN transcriptional control. Dysregulation of key chromatin modifiers such as *EZH2* and *DNMT3A* supports leukemic self-renewal and resistance, while SUMOylation influences DNA repair and transcription, emerging as a promising therapeutic target ^40, 41^. *PTEN*, a tumor suppressor often altered in AML, is highlighted here for its role in controlling PI3K-AKT signaling, with therapeutic implications for overcoming resistance and restoring apoptotic regulation ^42^.

Clusters 4, 6, and 7 reveal additional vulnerabilities. Cluster 4 (C4) features aberrant MAPK/ERK and PI3K/AKT signaling, key drivers of AML proliferation and chemoresistance, with added enrichment in immune receptor signaling pathways like FCGR-mediated phagocytosis. Cluster 6 (C6) centers on mitotic regulation and chromatid segregation, suggesting treatment-related disruption of chromosome stability. Cluster 7 (C7) is uniquely marked by mRNA regulation and TP53 signaling, with LS treatment appearing to restore TP53-related tumor-suppressive functions.

Collectively, the network analysis results reveal intricate therapeutic mechanisms employed by LS and RU treatments, underscoring both shared vulnerabilities, such as intracellular transport and epigenetic control, as well as distinct vulnerabilities, particularly in immune regulation and mitotic fidelity. Notably, this analysis demonstrates that *TP53, NPM1,* and *DNMT3A,* which rank among the most frequently mutated genes in AML patients, are uniquely targeted by the combinatorial therapies and not by the individual agents. This observation underscores the potential of LS and RU to exploit AML-specific genetic dependencies and to overcome compensatory pathways that often limit single-agent efficacy.

To further focus the molecular effects of combination-specific potential targets, we filtered enrichment terms in which there are genes targeted by each combination. For the LS regimen, we isolated nine genes exclusively modulated by LS, which are in four clusters and involved in different reactome pathways (Fig. 9A). Cluster three supplies the largest fraction of these LS-specific hits by contributing 4 genes. *RBBP7* alone is annotated to regulation of *PTEN* gene transcription and cellular senescence, underscoring the capacity of LS to re-engage tumor-suppressive *PTEN* programs and senescence barriers. This protein regulates cell proliferation by binding the retinoblastoma protein and recruiting histone-deacetylase complexes to repress cell-cycle genes; its dysregulation in AML is implicated in epigenetic reprogramming that promotes leukemic proliferation and differentiation blocks ^43,44^. The triad *DNMT3A, ASH2L,* and *RBBP7* are jointly enriched in epigenetic and chromatin regulation, revealing that LS uniquely engages epigenetic machinery, particularly DNA and *H3K4* methylation readers and writers, to recalibrate leukemic transcriptional programs. *DNMT3A*, a *de novo* DNA methyltransferase, is mutated in approximately 20–25% of AML cases, most often at R882, leading to altered CpG methylation landscapes, impaired hematopoietic stem-cell differentiation, and association with poor overall survival ^45^. *ASH2L*, a core member of MLL/SET1 H3K4 methyltransferase complexes, is overexpressed in FLT3-ITD–positive AML and supports leukemic cell viability, whereas *ASH2L* knockdown induces apoptosis, identifying it as an AML dependency ^46^. Finally, *DNMT3A* and BRCA1 map to SUMOylation, pointing to a combination-specific destabilization of SUMO-regulated chromatin modifiers and DNA repair complexes.

**Figure 9.**
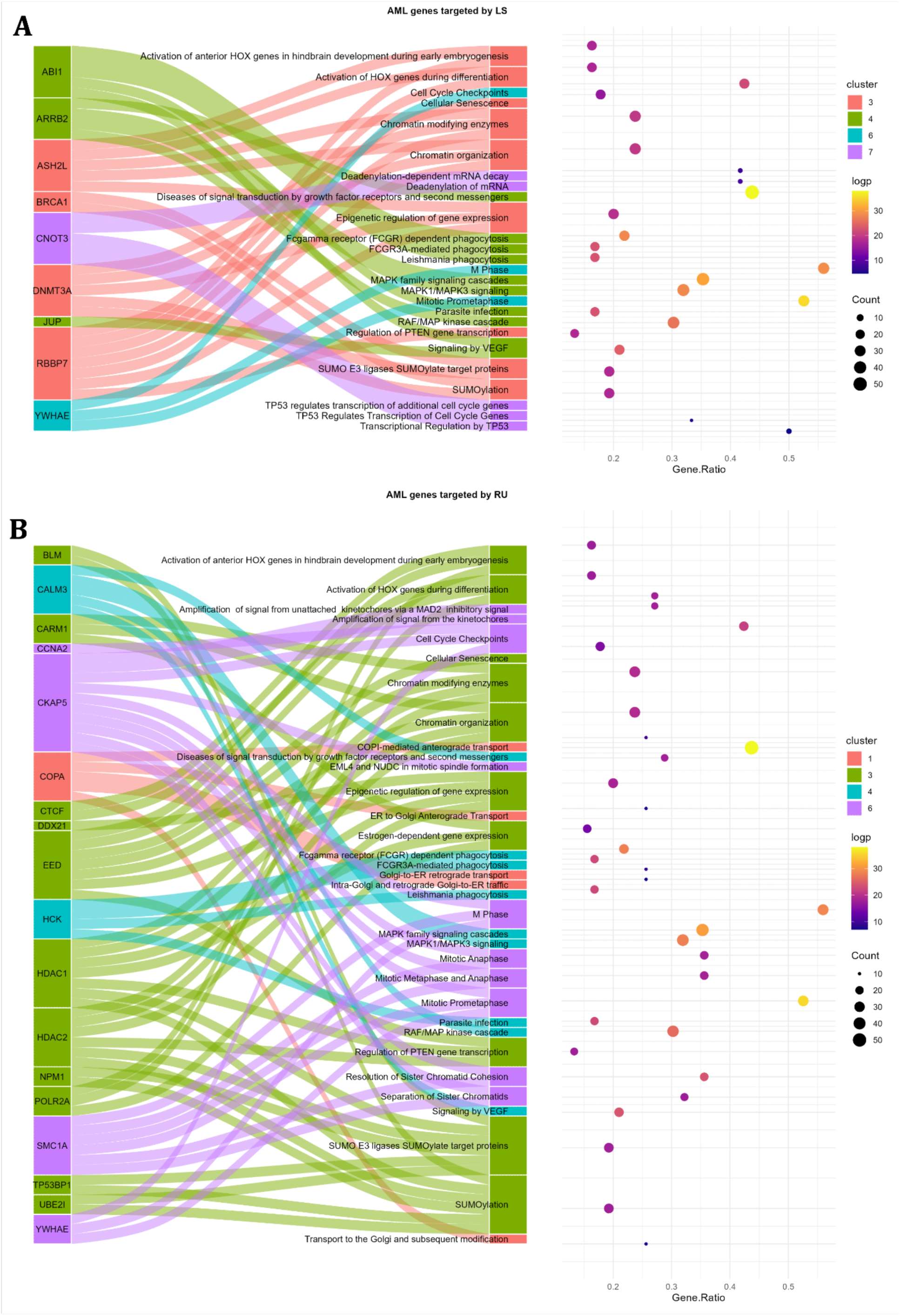
Pathway enrichment for LS- and RU-specific AML genes. (A) LS-specific genes: The Sankey plot (left) shows unique LS cluster genes (left nodes) and their associated enriched pathways (right nodes), with flows colored by cluster assignment. The adjacent dot plot (right) summarizes enrichment statistics for each pathway: the gene ratio (x-axis), number of genes (dot size), and significance (-log10 adjusted p-value, color). (B) RU-specific genes: As in (A), the Sankey plot and dot plot display RU cluster-specific genes and their enriched pathways. For both panels, pathway labels are identically ordered across plots to allow direct visual comparison.

C4 is represented in LS only by *JUP* and *ABI1*, which both feed into the signaling by *VEGF*. This suggests LS can implicate cell–cell adhesion and actin-remodeling hubs (*JUP*, junction plakoglobin; *ABI1*, Abl-interactor 1) in *VEGF* receptor trafficking or downstream signaling, potentially modulating the leukemic vascular niche ^47,48^. Moreover, *ARRB2* (β-arrestin 2), which scaffolds *MAPK1/MAPK3* (ERK1/2) activation, is upregulated in AML, sustaining ERK signaling that drives proliferation and survival. Hyperactivation of *MAPK1/MAPK3* is a hallmark of AML, transmitting growth factor and cytokine cues to support leukemic cell expansion and therapy resistance ^49^, so targeting this protein is highly effective in AML treatment.

In Cluster 6 (C6), the single protein *YWHAE* is mostly involved in the M Phase Regulation and Mitotic Chromatid Segregation. In AML, *YWHAE* silencing paradoxically increases proliferation, invasion, and migration through up-regulation of *CDC25B* and *MYC*^50^, and conversely, its engagement enhances p53-mediated apoptosis when combined with *HDAC* and *MDM2* inhibitors ^51^. *YWHAE* is in fusion with *NUTM2* and acts as a scaffold for *RAF/MEK/MAPK* activation, driving cyclin D1 up-regulation and increased proliferation ^52^.

Another noteworthy target in C7 is CNOT3, which stands out as the only LS-specific hit and is associated with mRNA regulation and TP53 signaling. The presence of *CNOT3* in this cluster highlights the dual mechanism of action of LS, destabilizing oncogenic transcripts through the CCR4–NOT deadenylase complex while reactivating p53-mediated cell cycle checkpoints ^53^. Together, these nine LS-specific perturbations carve out four major vulnerabilities in AML: (1) late-M-phase checkpoint control via *YWHAE*, (2) SUMO- and DNA-methylation–dependent epigenetic reprogramming *(RBBP7, DNMT3A, BRCA1, ASH2L)*, (3) VEGF pathway modulation through junctional and cytoskeletal adaptors *(JUP, ABI1)*, and (4) coupled mRNA destabilization and p53-axis reactivation via *CNOT3*.

By filtering our prioritized clusters for those genes uniquely modulated by RU, we isolated 18 RU-exclusive targets spanning four clusters (C1, C3, C4, and C6) (Fig. 9B) and these genes unmask combination-specific mechanisms. Considering C1, RU uniquely modulates *COPA*, a core *COPI* coatomer subunit, enriching four closely related pathways to intracellular transport and organelle trafficking. *COPA* mediates the retrieval of ER-resident proteins and SNAREs to maintain Golgi and ER homeostasis ^54^. The unique targeting of *COPA* by RU suggests that dysregulating secretory pathway logistics is an AML vulnerability exploited by this regimen.

Cluster 3 (C3), which is defined by epigenetic and SUMOylation processes, contributed the bulk of RU-specific genes such as *HDAC1, HDAC2, EED, CARM1, UBE2I, NPM1, BLM, TP53BP1,* and *CTCF,* thereby collapsing the histone deacetylase, Polycomb, and SUMO-mediated pathways that AML cells exploit for epigenetic plasticity. Critically, RU uniquely modulates *NPM1* and *TP53*: *NPM1* mutations occur in ∼30% of adult AML (and over 50% of normal-karyotype cases), producing a cytoplasmic *NPM1c* that disrupts ribosome biogenesis, centrosome duplication, genomic stability, and ARF–p53 tumor suppression to sustain leukemic transcriptional programs ^55,56^. *TP53* mutations, found in approximately 5– 10% of *de novo* AML, especially in elderly and therapy-related cases, abrogate DNA-damage checkpoints and apoptosis, driving chemo-resistance and dismal outcomes ^57^. By targeting SUMO ligation (*UBE2I*), chromatin boundary maintenance *(CTCF)*, and repair scaffolds *(TP53BP1)*, RU further dismantles the nucleolar integrity and genome-surveillance functions anchored by *NPM1* and *p53*, underscoring its capacity to erode the core epigenetic and checkpoint defenses in AML.

In Cluster 4 (C4), *CALM3* and *HCK* were the RU-exclusive hubs, linking calcium-regulated *MAPK1/MAPK3* and VEGF receptor signaling with FCGR-mediated phagocytosis pathways. This pattern implicates RU in concomitantly blocking proliferative MAPK/VEGF cues and disrupting AML–immune receptor crosstalk. Also worth noting, C6 yielded SMC1A and CKAP5 as unique mitotic regulators. By targeting cohesin dynamics and spindle assembly factors, RU appears to induce mitotic catastrophe through irreparable chromatid segregation errors. Together, these eleven RU-specific perturbations coalesce into four mechanistic vulnerabilities: intracellular trafficking disruption via *COPA*; epigenetic and SUMO network collapse through chromatin modifiers and DNA-repair regulators; a dual blockade of MAPK/immune receptor signaling by *CALM3* and *HCK*; and mitotic catastrophe induction via cohesin and spindle machinery.

This combination-focused view highlights how RU uniquely undermines AML cell survival by derailing logistics, rewriting the epigenome, rewiring intercellular signaling, and collapsing mitotic fidelity.

### PTM Enrichment Analysis of Differential Soluble Proteins

We were also interested in investigating the post-translational modification (PTM) status of protein targets in order to assess their potential role in therapy response. To get an idea of what PTMs might be potentially present on target proteins in extract and intact samples, we utilized the PEIMAN2 R package ^58^. Interestingly, significantly enriched PTM terms were different between extract and intact samples (**Supplementary** Figures 2 **& 3).** For example, phosphorylation of tyrosine, serine, and threonine together with acetylation of lysine showed significant enrichment in living cells. However, samples treated with drugs after extraction of proteins showed significant enrichment in dimethylation of arginine and acetylation of lysine, methionine, and alanine. These observations imply that acetylation and methylation may be involved in direct binding of drugs, while phosphorylation and acetylation are signatures of the secondary cell response to the treatment.

To confirm predictions of the PEIMAN2, we searched proposed PTMs in the proteomics data. Collectively, among 1280 protein targets after ruxolitinib, ulixertinib, and RU treatment in living cells, we found phosphorylation of tyrosine on 10 proteins (Fig. 10A). On the other hand, treatment with LS (either single drug or combination) in living cells yielded 950 proteins, among which 42 carried various PTMs (phosphorylation of serine and threonine, and acetylation of lysine) (Fig. 10B). In the case of extract samples, treatment with RU and LS showed 19 (975 in total) and 28 (626 in total) proteins with PTMs, respectively, comprising acetylation on alanine, methionine, and lysine, and dimethylation of arginine (**Supplementary File S4**). Overall, this data shows that there is an involvement of PTMs in a cell response to the anti-AML treatment.

**Figure 10:**
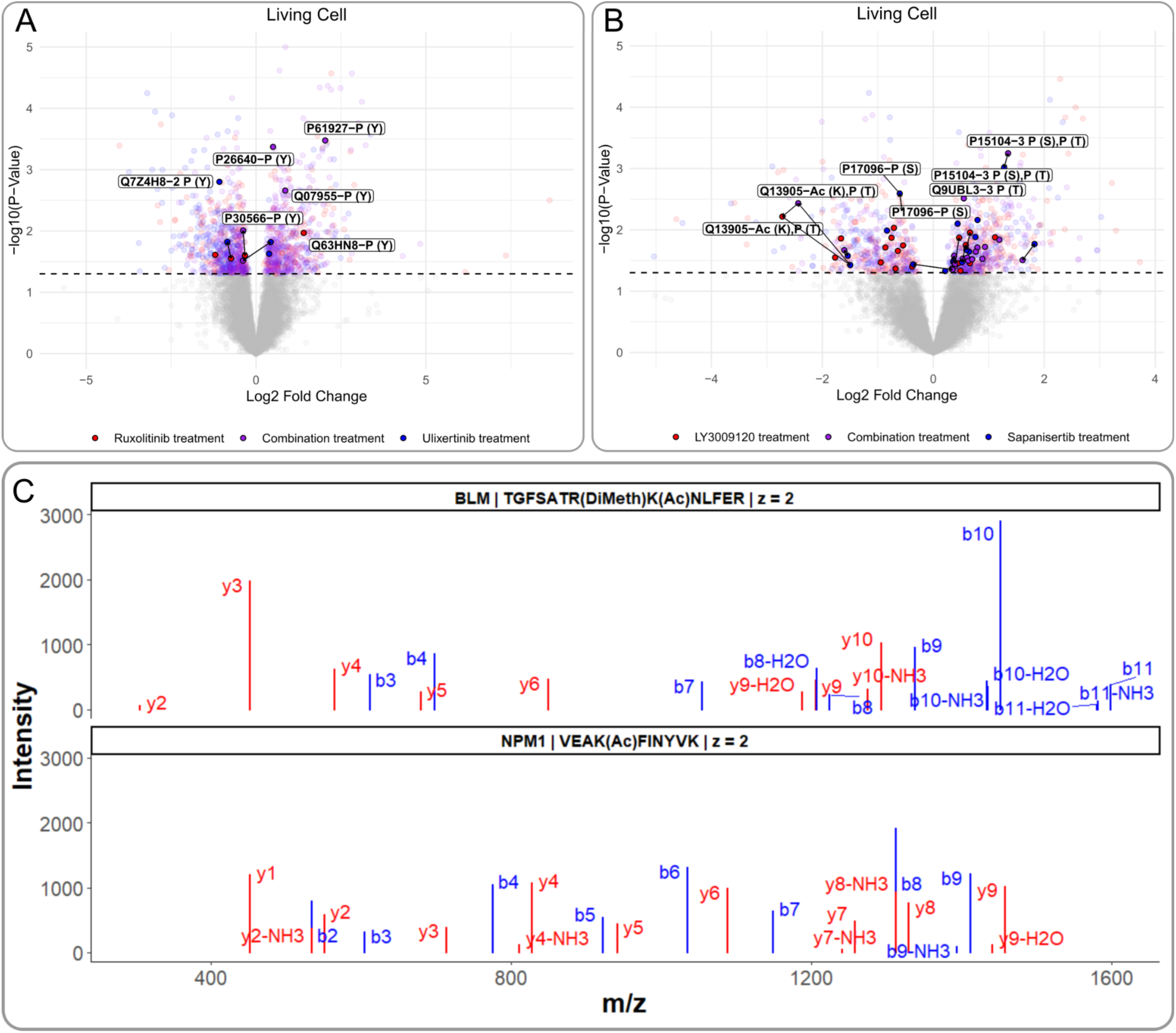
Protein targets of therapy and their PTM status in living cells treated with (A) ruxolitinib (3 uM) (red), ulixertinib (3 uM) (blue), and their combination, RU (purple); and (B) LY3009120 (red), sapanisertib (500 nM) (blue), and their combination, LS (purple). Highlighted points indicate proteins with PTM(s) identified in mass spectrometry data. The PTM(s) name and the site of modification are listed after the protein identifier through a hyphen. PTMs listed through commas were identified on the same peptide. Arrows indicate solubility change in the same protein from different treatments. P(Y)-tyrosine phosphorylation, P(T)-threonine phosphorylation, P(S)-serine phosphorylation, and Ac(K)-lysine acetylation. (C) MS/MS spectra for peptides with modifications from *BLM* and *NPM1* proteins. Only b-(blue) and y-fragment (red) ions are plotted for simplicity. The list of all CoPISA protein targets with PTMs is provided in a table as Supplementary File S4.

Next, we were interested in whether AML-related CoPISA protein targets (Fig. 10) are post-translationally modified. PTMs were detected on proteins across multiple clusters in both RU and LS treatments, with some overlap and distinct patterns between the two. Under RU treatment, *DYNC1I1* (cytoplasmic dynein 1 intermediate chain 1) from C1 was acetylated on K42, a modification not previously reported. *BLM* (RecQ-like DNA helicase *BLM*) and *NPM1* were also modified, with *BLM* showing dimethylation and both *BLM* and *NPM1* displaying acetylation (Fig. 12C). In addition to acetylation and dimethylation, tyrosine phosphorylation was observed; for example, *GSK3A/B* in C4 was phosphorylated on Y279, and *MCM2* in C6 on Y137. *PPIA* in C8 was also acetylated on K155. Under LS treatment, the highest number of modified protein targets was found in C3. *ASH2L* was phosphorylated simultaneously at T307, T310, and T311, while *CBX3* showed phosphorylation at S95 and S176. *H4C1* was acetylated at K9 and K13 on the same peptide. In contrast, C6 included only one modified protein, *YWHAE*, with N-acetylation on M1. In C7, *CNOT3* was phosphorylated on S299.

Surprisingly, several AML-related CoPISA protein targets that were modified were exclusively targeted by combination treatment (*AND* gate). For example, *BLM* and *NPM1* proteins are involved in the SUMOylation process as proteins that are SUMOylated and uniquely targeted by RU and not by any single drug. Notably, protein *BLM* had dimethylation (R483), which has not been reported before, and acetylation (K484) simultaneously on the same peptide. Additionally, on both proteins, lysines are sites for acetylation as well as SUMOylation (K484 in *BLM* and K267 in *NPM1*). Protein *NPM1* is shown to be retained in the nucleoplasm when acetylated, while its nucleolar residency is prevented by SUMOylation ^59,60^. It implies that different modifications on the same site can trigger different outcomes for the cell. *BLM* is a helicase that is involved in DNA replication and repair ^61^. While acetylation regulates the function of *BLM* in response to DNA damage, SUMOylation is crucial for interaction with *RAD51* protein in homologous recombination repair, localization to PML nuclear bodies, and *BLM* turnover ^62,63^. Thus, different functions and the localization of the *BLM* protein can be triggered through the same modification site.

These findings highlight the complex interplay between identified PTMs and AML-related protein function, suggesting that specific modifications induced by RU and LS combination treatments may act as key regulatory switches controlling protein localization, interactions, and activity in leukemia cells.

## Discussion

Combinatorial therapy has long been a cornerstone in oncology, particularly for aggressive malignancies like AML, due to its potential to overcome resistance and improve patient outcomes^64^. The classical rationale behind combinatorial regimens was to reduce the likelihood of resistance by employing multiple agents with distinct mechanisms of action, thus reducing the likelihood of clonal escape ^65^. This principle has yielded durable responses in certain hematologic malignancies; however, in AML, the substantial genetic and epigenetic heterogeneity among patients continues to pose formidable challenges for identifying effective and broadly applicable combinations ^66^. Moreover, the frequent emergence of therapy-resistant clones and the dose-limiting toxicities associated with multi-drug regimens have constrained their therapeutic window and limited their translational success^67^. These obstacles underscore the urgent need for strategies that rationally design and mechanistically dissect drug combinations to maximize efficacy while minimizing toxicity.

In this study, we introduced the CoPISA workflow (Proteome Integral Solubility/Stability Alteration Analysis for Combinations) to unravel the molecular underpinnings of combinatorial therapies in AML at a proteome-wide scale. Unlike traditional methods that focus on additive or synergistic effects of individual agents, CoPISA captures drug-induced changes in protein solubility that specifically arise from the combined action of drugs, providing mechanistic insights into cellular responses unattainable by monotherapies. Applying CoPISA to two rationally designed AML drug pairs ^13^, LY3009120-sapanisertib (LS) and ruxolitinib-ulixertinib (RU), we observed that these combinations elicit cooperative effects on protein stability, indicative of complex biological interactions rather than mere additive perturbations. These findings led us to propose the concept of *conjunctional targeting or inhibition*, describing a phenomenon where two drugs together interact and inhibit a protein target more effectively than either agent alone. This “AND-gate”-like behavior represents a conceptual advance over traditional models of synergy and aligns with emerging evidence that multi-agent therapies can reprogram cellular networks in fundamentally distinct ways

Previous proteomic profiling in AML has revealed subtype-specific vulnerabilities. For example, Jayavelu et al. demonstrated proteogenomic subtypes defined by mitochondrial activity, ribosome biogenesis, and proteostasis, which predict therapy response ^68^. Similarly, Tyner et al. identified signaling heterogeneity across AML samples using functional genomics, emphasizing MAPK and JAK/STAT pathways as potential combination targets ^5^. While these studies highlighted the diversity of AML dependencies, they did not interrogate emergent proteomic changes unique to drug combinations. Our CoPISA workflow advances the field by resolving combination-specific protein stability shifts, enabling mechanistic dissection of cooperative drug effects at a proteome-wide scale. Furthermore, classical synergy screens, such as those employing Bliss ^69^ or Loewe ^70^ models, quantify functional synergy but offer limited mechanistic resolution. In contrast, CoPISA directly captures the conformational and stability changes of proteins in response to drug co-treatment, providing insights into network rewiring and actionable vulnerabilities. Notably, our finding that 47% of LS-perturbed and 54% of RU-perturbed AML-associated genes were unique to the combinations highlights the prevalence of conjunctional effects in AML cells.

The integration of *AND* and *OR* logic gates into our CoPISA analysis provides a powerful systems-level perspective on how LS and RU combinations reprogram leukemic cell function to achieve synergistic effects. The enrichment of SUMOylation, chromatin condensation, and mitotic control pathways in the LS *AND* gate highlights the ability of this combination to destabilize genome integrity and impair cell division fidelity, two hallmarks of AML proliferation and resistance ^71^. Simultaneously, disruption of RNA processing and immune-related pathways in intact cells suggests that LS not only targets intrinsic leukemic signaling but also modulates the tumor microenvironment to potentially enhance immune clearance ^72^. The complementary effects of RU, which prominently engages transcriptional regulation, epigenetic modulation, and apoptosis pathways under the *AND* gate, underscore its capacity to restore tumor suppressor networks and induce metabolic collapse in AML cells ^73,74^. Notably, the convergence of LS and RU effects on phospholipid metabolism and RNA splicing across both gating models and cellular fractions suggests that these core biosynthetic processes represent shared vulnerabilities in AML. Together, these findings provide a mechanistic rationale for how conjunctional targeting in LS and RU treatments exerts multi-axis pressure on AML survival networks, overwhelming adaptive resistance and supporting the development of rational, precision-guided combinatorial regimens.

The functional network analysis of AML-associated genes and PTM enrichment analysis provide critical mechanistic insight into how the LS and RU combinations reprogram leukemia-specific signaling and survival pathways to achieve therapeutic synergy. LS treatment prominently engaged modules involved in epigenetic regulation *(DNMT3A, ASH2L, RBBP7)* and PTEN transcriptional control, suggesting a capacity to disrupt the chromatin architecture of AML and restore tumor-suppressive pathways such as PI3K-AKT inhibition ^75,76^. In contrast, RU targeted an overlapping yet distinct set of vulnerabilities, including *NPM1* and *TP53*, two of the most frequently mutated and therapeutically challenging nodes in AML. *NPM1* mutations, which occur in approximately 30% of AML cases, promote aberrant cytoplasmic localization and impair genomic stability and ribosomal biogenesis^77^. *TP53* mutations, present in ∼10% of AML and associated with chemoresistance and poor prognosis, disrupt DNA damage checkpoints and apoptosis ^78^. The engagement of RU with SUMOylation networks and cohesion/spindle assembly proteins (e.g., *SMC1A*, *CKAP5*), indicating a potential to induce mitotic catastrophe and disrupt genomic surveillance mechanisms essential for leukemic persistence ^79^. Shared perturbations in MAPK/ERK and PI3K/AKT signaling across both combinations highlight these pathways as central hubs of AML pathogenesis and promising points for combinatorial intervention ^80^. Collectively, these findings reinforce that LS and RU act not merely through additive inhibition but by simultaneously destabilizing multiple survival axes, including epigenetic, metabolic, and immune regulatory networks, to overwhelm AML cells’ adaptive capacity. This network-centric perspective also highlights potential biomarkers and actionable targets (e.g., *DNMT3A, NPM1, PTEN*) that could inform precision therapy approaches for genetically diverse AML subtypes. Finally, prospective clinical evaluation of LS and RU regimens, guided by biomarkers like *DNM3A*, *NPM1*, and *TP53* status, could advance precision medicine approaches for genetically diverse AML subtypes.

## Materials and Methods

### Ab Initio Simulation

The simulation of sigmoidal curves of 10000 hypothetical proteins was carried out using R, and melting temperatures *T*_m_, selected randomly in the range from 48 to 56°C. *N_t_* = 15 temperature points between 45 and 59°C with a 1°C step were chosen. Sigmoidal melting curves were simulated by calculating the relative intensity *I*(*T*) for a given temperature *T* as *Intensity*(*T*)*_control_* = *0.5 erf*[(*T_m_* − *T*)/*sqrt*(*T_m_*)/*2* where *erf* is the error function and *sqrt* is the square root function. Similarly, sigmoidal melting curves for treated samples was simulated as *Intensity*(*T*)*_treated_* = *0.5 erf*[(*T_m_* + Δ*T_m_* − *T*)/*sqrt*(*T_m_* + Δ*T_m_*)/*2*. Δ*T_m_* were simulated as random values in the range between −*3*°C and +*3*°C. The sum of measured signals for control and treated samples is calculated as *S_m_* = ∑_T=1_^15^ *Intensity*(*T*), for all temperature points. The treatment here was simulated for two hypothetical drugs to calculate *S_m_* values, i.e. *S_m_^control^*, *S_m_^1^* and *S_m_^2^*. In the next step, we calculated the two Δ*S_m_* for both hypothetical drugs, i.e., Δ*S_m_^1^* and Δ*S_m_^2^* in 10,000 proteins to depict versus each other.

### Cell culture

MOLM-13, MOLM-16, SKM, and NOMO-1 cells were obtained from Deutsche Sammlung von Mikroorganismen und Zellkulturen (DSMZ, Germany) and cultured in RPMI-1640 medium (Gibco, Thermo Fisher Scientific, USA) supplemented with 10–20% fetal bovine serum (FBS) (Gibco, Thermo Fisher Scientific), 2 mM L-glutamine (Lonza), and 100 units/mL penicillin/streptomycin (Gibco, Thermo Fisher Scientific) at 37°C and 5% CO_2_. Cells were harvested and centrifuged at 400 g for 4 min.

### Cell lysate-based CoPISA

Cells were cultured until they reached the desired density of 2 × 10⁶ cells per treatment with seven different treatment groups for each cell line, performed in triplicate. A 24 ml volume of cell culture was pelleted by centrifugation at 600 g for 5 minutes, followed by two washes with phosphate-buffered saline (PBS). After a final pelleting, cells were resuspended in PBS (GIBCO) supplemented with protease inhibitors (Roche, Switzerland). The cell mixture was lysed through 4 cycles of freezing in liquid N₂ and then thawing at 35°C. At the final step, cells were cleared by centrifugation at 10,000 g for 10 min at 4°C. Samples were then treated with single or combination drug regimens at the following concentrations: LY3009120 500 nM, sapanisertib 500 nM, ruxolitinib 3 μM, and ulixertinib 3 μM for 15 min at room temperature. For the treatment of control samples, 0.1% DMSO was used. After treatment, the samples were aliquoted into 12 equal volumes for subsequent heat treatments. Each aliquot was subjected to a 3-minute heat treatment at temperatures ranging from 48°C to 59°C in 1°C intervals using a thermocycler (Applied Biosystems Veriti). After equilibrating at room temperature for 3 min, samples from different temperatures for each drug treatment were pooled and subjected to ultracentrifugation at 100,000 g for 20 minutes at 4°C (Beckman Coulter Optima Max-XP). The soluble proteins from the supernatant were collected for further analysis.

### Living cell-based CoPISA

Cells (2 × 106 cells per condition) were treated with a single or combination drug with concentrations mentioned in 6-well plates and incubated at 37°C with 5% CO₂ for 60 min. Following treatment, the cells were washed twice with PBS and resuspended in 320 µL of PBS supplemented with a protease inhibitor. An equal volume of each sample was transferred to the PCR plate and was heated for 3 min at the same temperatures used in the lysate-based protocol. The samples were allowed to cool down at room temperature for 3 min and then applied for four cycles of freezing cells in liquid N₂ and thawing at 35°C. After pooling the samples from different temperature points for each treatment, the soluble proteins were isolated via ultracentrifugation at 100,000 g for 20 min at 4°C. At this point, samples can be flash frozen and stored at −80°C for future analysis.

### Proteomics Sample Processing

Protein concentration was measured using the PierceBCA Protein Assay Kit (Thermo), and the volume corresponding to 27 μg of protein was transferred from each sample to prepared filter tubes (Amicon Ultra-0.5 mL Centrifugal Filters 10KD). The volume of each sample was adjusted to an equal level using 20 mM 4-(2-hydroxyethyl)-1-piperazinepropanesulfonic acid (EPPS) buffer, then was reduced in 10 mM dithiothreitol (55°C, 30 min) and alkylated in 50 mM iodoacetamide (25°C, 60 min in darkness). After reduction and alkylation, samples were centrifuged for 20 min at 14000 g, followed by two washes. EPPS buffer was then added to the Amicon filter tubes containing samples. Proteins were digested overnight at 37°C with vigorous shaking by trypsin at a ratio of 1:50. The resulting peptides were collected by centrifugation and rinsing the filter tubes with EPPS. Prior to labeling, equal amounts of each sample from the different treatment groups were mixed for separate labeling. A total of 25 μg from each sample was labeled using tandem mass tags (TMT) according to the manufacturer’s protocol (Thermo Fisher Scientific). Three technical replicates for each treatment (drug A, drug B, AB, and control), along with one mixed sample of each condition, were pooled together, resulting in a final mixture of 325 μg from 13 samples.

### Peptide fractionation and LC MS/MS analysis

TMT-labeled peptides were fractionated by high-pH reversed-phase separation using an XBridge Peptide BEH C18 column (3.5 µm, 130 Å, 1 mm x 150 mm, Waters) on an Ultimate 3000 system (Thermo Scientific). Peptides were loaded on the column in solution A, and the system was run at a flow of 42 µl/min. The following gradient was used for peptide separation: from 2% B to 15% B over 3 min to 45% B over 59 min to 80% B over 3 min, followed by 9 min at 80% B, then back to 2% B over 1 min, followed by 15 min at 2% B. Solution A was 20 mM ammonium formate in water, pH 10, and Solution B was 20 mM ammonium formate in 90% acetonitrile, pH 10. Elution of peptides was monitored with a UV detector (205 nm, 214 nm), and a total of 36 fractions were collected, pooled into 12 fractions using a post-concatenation strategy as previously described ^81^, and dried under vacuum.

Dried peptides were resuspended in 0.1% aqueous formic acid and subjected to LC–MS/MS analysis using an Orbitrap Eclipse Tribrid Mass Spectrometer fitted with an Ultimate 3000 nano system and a FAIMS Pro interface (all Thermo Fisher Scientific) and a custom-made column heater set to 60°C. Peptides were resolved using an RP-HPLC column (75 μm × 30 cm) packed in-house with C18 resin (ReproSil-Pur C18–AQ, 1.9 μm resin; Dr. Maisch GmbH) at a flow rate of 0.3 μl/min. The following gradient was used for peptide separation: from 2% B to 12% B over 5 min to 30% B over 70 min to 50% B over 15 min to 95% B over 2 min, followed by 18 min at 95% B, then back to 2% B over 2 min, followed by 18 min at 2% B. Solution A was 0.1% formic acid in water, and Solution B was 80% acetonitrile and 0.1% formic acid in water.

The mass spectrometer was operated in DDA mode with a cycle time of 3 s between master scans. Throughout each acquisition, the FAIMS Pro interface switched between CVs of −40 V and −70 V with cycle times of 1.5 s and 1.5 s, respectively. MS1 spectra were acquired in the Orbitrap at a resolution of 120,000 and a scan range of 400 to 1600 m/z, AGC target set to “Standard” and the maximum injection time set to “Auto”. Precursors were filtered with a precursor selection range set to 400–1600 m/z, monoisotopic peak determination set to “Peptide”, charge state set to 2 to 6, a dynamic exclusion of 45 s, a precursor fit of 50% in a window of 0.7 m/z, and an intensity threshold of 5e3. Precursors selected for MS2 analysis were isolated in the quadrupole with a 0.7 m/z window and collected for a maximum injection time of 35 ms with the AGC target set to “Standard”. Fragmentation was performed with a CID collision energy of 30%, and MS2 spectra were acquired in the IT at a scan rate of “Turbo”.

MS2 spectra were subjected to real-time search (RTS) using a human database containing 20362 entries downloaded from Uniprot on 20200417 using the following settings: Enzyme was set to “Trypsin”, TMTpro16plex (K and N-term) and Carbamidomethyl (C) were set as fixed modifications, Oxidation (M) was set as a variable modification, maximum missed cleavages were set to 1, and maximum variable modifications to 2. Maximum search time was set to 100 ms, the scoring threshold was set to 1.4 XCorr, 0.1 dCn, 10 ppm precursor tolerance, charge state 2, and “TMT SPS MS3 Mode” was enabled. Subsequently, spectra were filtered with a precursor selection range filter of 400–1600 m/z, precursor ion exclusion set to 25 ppm low and 25 ppm high, and isobaric tag loss exclusion set to “TMTpro”. Tandem MS product ions of precursors identified via RTS were isolated for an MS3 scan using the quadrupole with a 2 m/z window, and ions were collected for a maximum injection time of 200 ms with a normalized AGC target set to 200%. SPS was activated, and the number of SPS precursors was set to 10. Isolated fragments were fragmented with normalized HCD collision energy set to 55%, and MS3 spectra were acquired in the Orbitrap with a resolution of 50,000 and a scan range of 100 to 500 m/z.

The acquired raw files were analyzed using the MaxQuant software (v2.6.5.0). Spectra were searched against a human protein sequences database (downloaded from Uniprot on 20241010). Group-specific parameters were used as follows. The type of the search was set to Reporter MS3. Isobaric labels and their correction factors were provided according to the manufacturer kit sheet. Normalization was set to a weighted ratio to the reference channel. Variable modifications were set to oxidation of methionines and acetylation of N-termini, while fixed modification settings contained only carbamidomethylation of cysteines. The maximum number of modifications per peptide was set to 3. Two missed cleavages with Trypsin/P were allowed. For the refined search, all settings were kept as in the original with the addition of variable modifications as proposed by the PEIMAN2 R package, with five as a maximum number of modifications per peptide.

### LC-MS Analysis of Drug Combinations

Ten picomoles from each drug, separately or combined, were injected into LCMS (Eksigent Ekspert 400 coupled to TripleTOF 6600 (Sciex, USA)). These analytes were separated using a linear gradient of 15 min comprising 8 min from 3% to 35% of solution B (0.1% formic acid/acetonitrile) and 2 min to 90% of solution B on a reverse phase C4 column (0.3 x 20 mm, Phenomenex, USA). The solution A was 0.1% formic acid, and the flow rate was at 5 µl/min. The mass spectrometer was set in TOF MS mode with the following parameters: Ion spray voltage was at 5500 volts, and ion source and curtain gases were 10 and 30 liters per minute, respectively. TOF MS was acquired from m/z 200 to 1500. Following LC-MS acquisition, raw files were analyzed by PeakView^®^ software, version 2.2 (Sciex). Drug ions were extracted using their corresponding m/z +/- 0.5 Da.

### PTM Enrichment Analysis

PTM enrichment analysis was performed utilizing the PEIMAN2 R package. Firstly, a ranked list (by p-value of statistical test from the limma package) of differentially soluble proteins with their UniProtAC was created for every treatment condition. Next, the function runPSEA() was executed to perform protein set enrichment analysis. Results were filtered based on the number of proteins enriched for every PTM term (≥10) and significance of the enrichment (0.05). These PTMs were provided as modification values as input to perform the refined search by MaxQuant proteomics software.

### Statistical Analysis

Proteomics data were obtained from primary Peptide Spectrum Match (PSM) reports, providing raw intensity values of peptides identified across experimental conditions. Invalid or missing treatments were excluded, and intensity values were normalized to account for differences in protein abundance across samples. A pooled control sample was used for further normalization, and protein intensities were averaged across replicates to create matrices for each treatment condition.

Linear modeling with empirical Bayes adjustments was applied for statistical analysis. Comparisons were made between control (C) and drug treatment conditions (R, U, RU, L, S, LS). Contrast matrices were defined to account for single and combination drug effects, with significance thresholds set at p-value < 0.05. Volcano plots were generated for each treatment comparison to visualize significantly altered proteins based on logFC and p-values. Fold change plots comparing each combination of treatments (comb vs. drug 1 and comb vs. drug 2) were generated, with primary plots displaying fold changes across the full dataset range and inset plots highlighting a focused subset of the most significant proteins based on p-values.

Due to the complexity of comparing single drugs and combinations in the CoPISA method and the lack of an a *priori* error model in a low-power proteomics context^82^, direct multiple hypothesis correction was not feasible. To address this, we tested our candidate selection procedure and estimated the false discovery rate (FDR) for proteins with significant solubility shifts. This involved permuting the protein S_m_ values across replicates within the dataset to generate 1000 randomized permutations ^83–85^. The same selection criteria were applied to these permuted datasets as to the original data, enabling us to evaluate the robustness of our approach and ensure confidence in the identified candidates.

### AML-Related Network Construction and Functional Annotation Workflow

The signaling network dataset was obtained from Reactome and preprocessed by removing duplicate edges. Subsequently, AML-associated gene mutations were downloaded from two sources: DEPMAP (for cell lines) and the FIMM-AML cohort (for patient data). Mutations were filtered using a stringent threshold (p-value < 10e-8), and only those classified as high-impact, such as frameshift mutations, nonsense mutations (introducing stop codons), and similar functionally disruptive variants, were retained. A directed network was then reconstructed by first filtering the global Reactome network to retain only the curated AML-mutated genes and known AML-related drug targets. This step ensured that only AML-associated nodes were included. For every pair of these AML-associated nodes, we computed the directed shortest path. All intermediate nodes appearing in any of these shortest paths were then used to induce a subnetwork from the original Reactome signaling network. The resulting edge list was extracted to form the final AML-specific signaling network. Network topology analysis was performed, and downstream analyses focused on the largest connected component, which comprised 4,626 nodes. Community detection was carried out using multiple algorithms, including Louvain, Walktrap, and Infomap. Among these, the Walktrap algorithm was selected for further analysis due to its superior modularity score. Following community detection, functional annotation was performed to identify biological pathways enriched in each cluster and treatment group. Proteins were grouped by treatment and cluster and mapped to Entrez IDs. Reactome pathway enrichment analysis was then carried out using ReactomePA with adjusted thresholds (p-value < 0.05, q-value < 0.2). This analysis highlighted key signaling pathways associated with AML-specific communities and drug responses within the reconstructed network.

## Contributions

M.J. conceived the study and designed the research questions. M.J. and A.A. provided scientific direction, supervised the project, and contributed to the manuscript preparation. E.G. led the experimental analyses and data interpretation, while E.Z. led the computational analyses and data visualization. U.V. participated in proteomics data analysis, and D.R. conducted the proteomics mass spectrometry experiments and data analysis. J.J.M. assisted with protein extraction and interpretation of signaling networks. R.S. and M.B. contributed to small-molecule mass spectrometry analysis. M.W. supported the interpretation of findings and the validation of the experimental design. C.A.H. provided cell lines and contributed to the interpretation of findings. E.K. and P.A.H. offered valuable inputs on data interpretation and visualization.

## Data availability

The mass spectrometry proteomics data have been deposited to the ProteomeXchange Consortium^86^ via the PRIDE ^87,88^ partner repository with the dataset identifier PXD066812.

## Supplementary

**Supplementary Figure 1.** U**p**set **plot showing AML gene overlaps across CoPISA treatment conditions.** Rows denote treatments; bar plots indicate the number of genes perturbed per treatment. Connected dots represent intersections between treatments, while bar heights above indicate intersection sizes. Colors map to individual treatments for visual clarity. This visualization highlights shared and unique AML gene responses, providing insights into convergent and divergent mechanisms of action.

**Supplementary Figure 2.** B**u**bble **plots from PEIMAN2 enrichment for all treatment comparisons in living cells:** (a) treatment with drug L; (b) treatment with drug S; (c) treatment with combination LS; (d) treatment with drug R; (e) treatment with drug U; (f) treatment with combination RU. The red circles with PTM highlighted in red denote the selected PTMs for the refined MaxQuant search.

**Supplementary Figure 3.** P**r**otein **targets of therapy and their PTM status in cell lysates** treated with (a) ruxolitinib (3 μM) (red), ulixertinib (3 μM) (blue), and their combination, RU (purple); and (b) LY3009120 (red), sapanisertib (500 nM) (blue), and their combination, LS (purple). Highlighted points indicate proteins with PTM(s) identified in mass spectrometry data. The PTM(s) name and the site of modification are listed after the protein identifier through a hyphen. PTMs listed through commas were identified on the same peptide. Arrows indicate solubility change in the same protein from different treatments. P(Y)-tyrosine phosphorylation, P(T)-threonine phosphorylation, P(S)-serine phosphorylation, and Ac(K)-lysine acetylation.

**Supplementary file S1:** Excel workbook comprising 30 worksheets, each named according to the group indicating IN (intact cell), EXT (extract samples), or IN_EXT (shared between intact and extract), and Subset denotes the treatment comparisons: R, U, RU, AND_RU, and OR_RU for the RU analysis, and L, S, LS, AND_LS, and OR_LS for the LS analysis. Each worksheet contains nine columns: UniProt accession (uniprot), log₂ fold-change (logFC), average expression (AveExpr), moderated t-statistic (t), raw p-value (P.Value), Benjamini–Hochberg adjusted p-value (adj.P.Val), log-odds of differential expression (B), HGNC gene symbol (gene_symbol), and Entrez Gene identifier (entrez_ids), reporting results for all genes in the corresponding set. “AND_…” sheets represent combination-specific targets, while “OR_…” sheets represent their unions.

**Supplementary file S2:** Excel file containing pathway enrichment analysis results for the Reactome database, organized into 12 sheets. Each sheet presents the enriched pathways identified for AND and OR gates and treatments under two experimental conditions: living cells and lysate extract. Analyses are separated for LS and RU combinations, providing detailed pathway insights for both treatments. Each sheet contains pathway names, reaction identifiers, statistical values, etc., supporting comprehensive comparison across conditions and combinations.

**Supplementary file S3:** Excel file presenting Reactome pathway enrichment for prioritized Walktrap-defined community in our AML signaling subnetwork, separated by treatment (LS and RU). For each cluster, it lists the Reactome pathway ID and name, the ratio of cluster genes annotated to that pathway versus background (GeneRatio/BgRatio), raw and FDR-adjusted P-values (pvalue/p.adjust/qvalue), the count of matching genes, a manual thematic label (mains), and the specific gene IDs driving enrichment. Additional columns (“overlap_L,” “overlap_S,” and “overlap_LS”) indicate whether the solubility shift of each protein was detected in each treatment, and columns (“L_only,” “S_only,” and “LS_only”) represent proteins unique to each treatment.

**Supplementary file S4:** Excel file presenting all identified proteins with PTM after doing the refined search based on the prediction of PEIMAN2. This file contains information about the treatments, peptide sequence, and modification sites.

## Acknowledgements

The authors acknowledge the Meilahti Clinical Proteomics Core Facility for Mass Spec. sample analysis (supported by HiLIFE and Biocenter Finland). We also thank Krister Wennerberg for helpful discussions and feedback on the manuscript. This study was financially supported by the Research Council of Finland [Grant 332454 to M.J.] and the Jane and Aatos Erkko Foundation [Grant 220031 to M.J.].

