## Supplementary Figures for "Shifting Beyond Classical Drug Synergy in Combinatorial Therapy through Solubility Alterations"

(4) [Technical University of Munich](https://scholar.google.com/citations?view_op=view_org&hl=en&org=1384714062669582586), Munich, Germany.

**Figures**

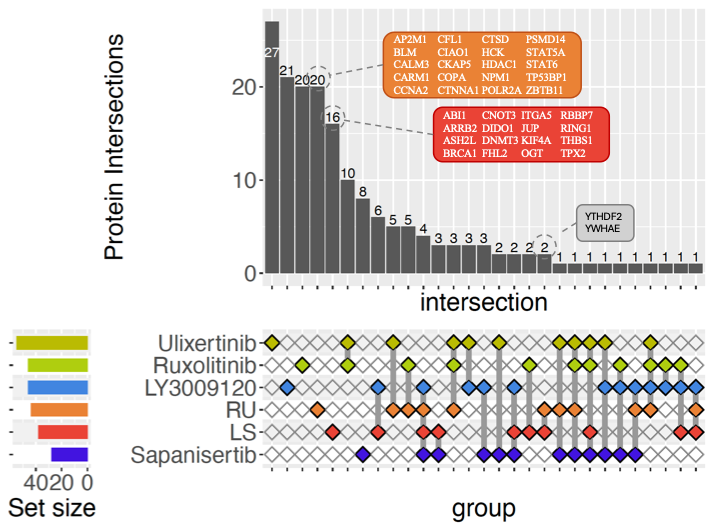

Figure 1. Upset plot showing AML gene overlaps across CoPISA treatment conditions. Rows denote treatments; bar plots indicate the number of genes perturbed per treatment. Connected dots represent intersections between treatments, while bar heights above indicate intersection sizes. Colors map to individual treatments for visual clarity. This visualization highlights shared and unique AML gene responses, providing insights into convergent and divergent mechanisms of action.

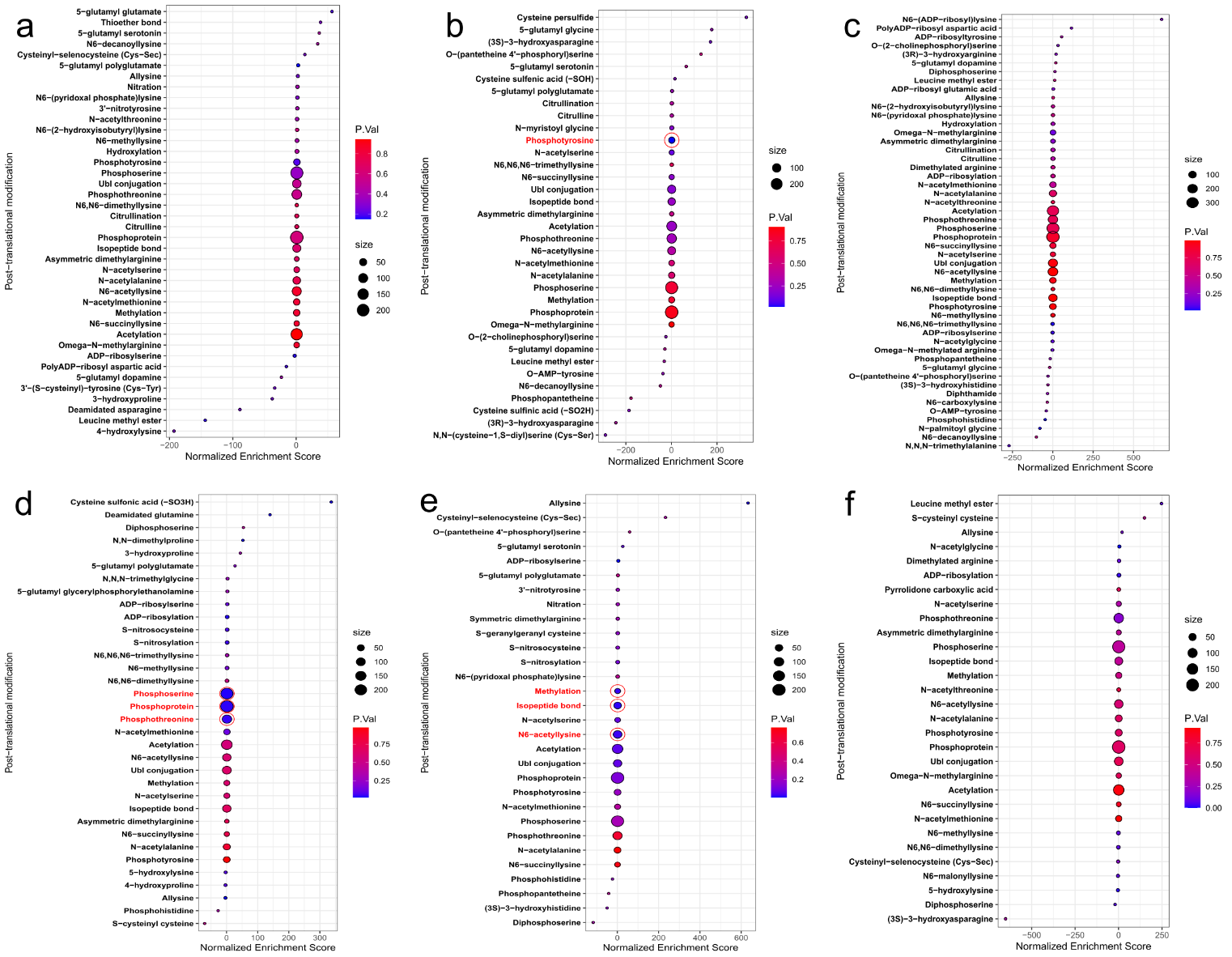

Figure 2. Bubble plots from PEIMAN2 enrichment for all treatment comparisons in living cells: (a) treatment with drug L; (b) treatment with drug S; (c) treatment with combination LS; (d) treatment with drug R; (e) treatment with drug U; (f) treatment with combination RU. The red circles with PTM highlighted in red denote the selected PTMs for the refined MaxQuant search.

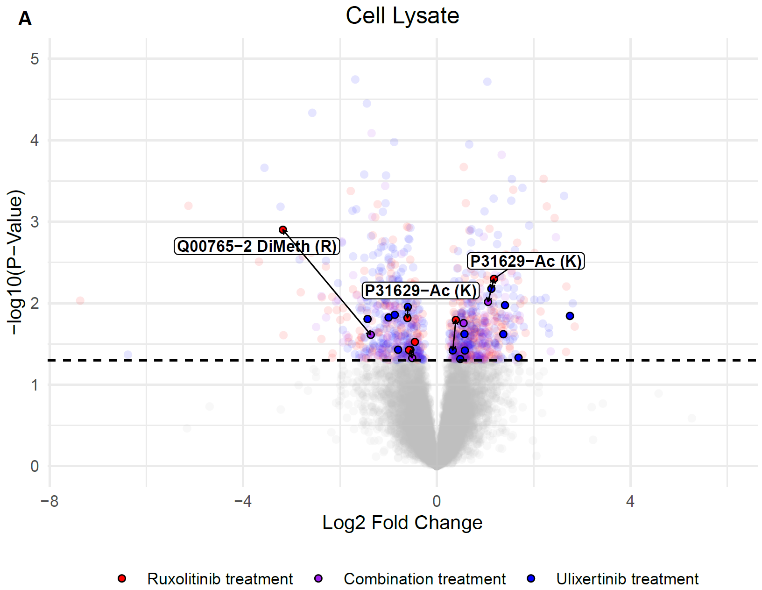

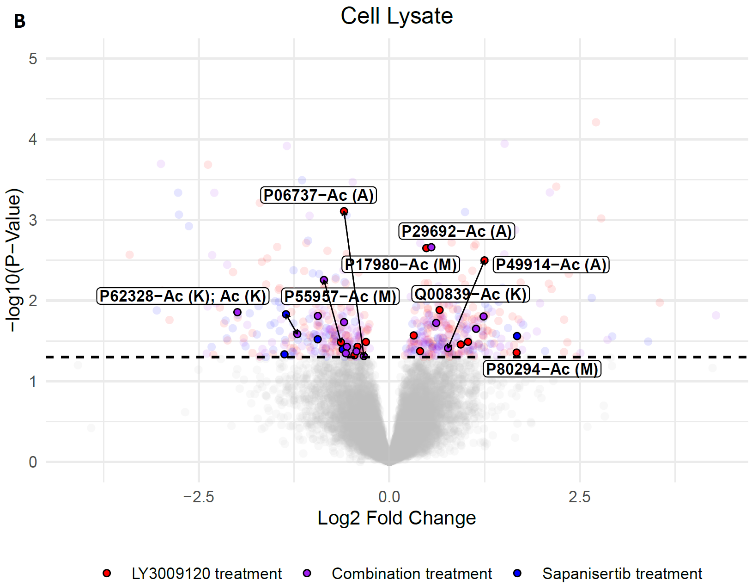

Figure 3. Protein targets of therapy and their PTM status in cell lysates treated with (a) ruxolitinib (3 μM) (red), ulixertinib (3 μM) (blue), and their combination, RU (purple); and (b) LY3009120 (red), sapanisertib (500 nM) (blue), and their combination, LS (purple). Highlighted points indicate proteins with PTM(s) identified in mass spectrometry data. The PTM(s) name and the site of modification are listed after the protein identifier through a hyphen. PTMs listed through commas were identified on the same peptide. Arrows indicate solubility change in the same protein from different treatments. P(Y)-tyrosine phosphorylation, P(T)-threonine phosphorylation, P(S)-serine phosphorylation, and Ac(K)-lysine acetylation.
